# Nucleotide and metalloid-driven conformational changes in the arsenite efflux ATPase ArsA

**DOI:** 10.1101/2025.03.21.644500

**Authors:** Shivansh Mahajan, Ashley E. Pall, Yancheng E. Li, Timothy L. Stemmler, Douglas C. Rees, William M. Clemons

**Author notes:** To whom correspondence should be addressed: Douglas C. Rees and William M. Clemons, Jr. **Email:** and. Timothy L. Stemmler: Medical University of South Carolina, Charleston, SC 29425.

## Abstract

A common mechanism of arsenic detoxification in bacteria is arsenite (As^III^) efflux facilitated by the ArsAB pump that couples metalloid transport to ATP hydrolysis. The cytoplasmic ATPase component, ArsA, binds and hydrolyzes ATP and facilitates the transfer of As^III^ to the integral membrane transporter, ArsB. The underlying molecular mechanism of As^III^ efflux by ArsAB remains unclear. ArsA is a member of the Intradimeric Walker A (IWA) family of ATPases that undergo dramatic nucleotide-dependent conformational changes to facilitate their respective biological functions. Similar conformational transitions in ArsA have been postulated to drive As^III^ binding and transport via ArsB but have not been demonstrated. Here, we report multiple structures of ArsA determined by single-particle cryogenic electron microscopy in an *open* MgADP-bound state, *open* MgATP-bound state, and a distinct *closed* MgATP-bound state liganded to As^III^. Using X-ray absorption spectroscopy, we confirmed that As^III^ coordinates three conserved cysteines at the metalloid-binding site of the *closed* state in a three-coordinate fashion. Coupled with biochemical characterization, our cryo-EM structures reveal key conformational changes in the ArsA catalytic cycle consistent with other members of the IWA family and provide the structural basis for allosteric activation of nucleotide hydrolysis by As^III^. This work enhances our understanding of how the ArsA catalytic cycle regulates metalloid efflux by ArsB.

## Introduction

With a natural abundance of 1.5-2 ppm, arsenic is a toxic metalloid that contaminates soil and groundwater, posing a widespread public health risk across the globe^1,2^. Several bacteria and archaea resist trivalent arsenite (As^III^) toxicity by using the arsenite efflux machinery encoded in the *ars* operon^3–6^. A major component of the *ars* operon is an integral membrane transporter, ArsB, that exports As^III^ out of the cell. Though ArsB can function alone, it often associates with a cytoplasmic ATPase, ArsA, that sequesters As^III^, and enhances the efficiency of As^III^ efflux via ArsB by coupling this process to ATP hydrolysis^7,8^. Intriguingly, properties of this ArsAB system are distinct from other ATP-dependent transporters, including ABC transporters and P-type ATPases, both in terms of the protein fold of the ATPase domain and that the ATPase activity is only required to improve the efficiency of transport^9–11^. In addition to the questions concerning the mechanism of toxic metalloid detoxification, the ArsAB transporter has garnered interest due to its biotechnological potential for arsenic bioremediation of contaminated environments^12,13^.

ArsA is a 63 kDa pseudodimeric ATPase that consists of two homologous nucleotide-binding domains covalently linked by a flexible ∼25-residue linker^14,15^ (Fig. S1A). The metalloid-binding site, composed of three conserved cysteines^16^, is located at the pseudodimer interface and can ligand As^III^ or antimonite (Sb^III^). Both domains of ArsA, namely the N- and C-domains, can support the binding and hydrolysis of ATP; however, the N-domain does so at a significantly higher rate than the C-domain^17^. ArsA belongs to a family of dimeric ATPases of diverse biological functions, known as the ‘Intradimeric Walker A (IWA) ATPases’ that are characterized by a conserved X-K-G-G-X-G-K-[T/S] Walker A or phosphate-binding loop (P-loop) motif^18,19^ (Fig. S1B). The first conserved Lys residue, referred to as the ‘IWA lysine’, is critical for facilitating nucleotide hydrolysis in these enzymes^20,21^. Nucleotide-dependent conformational changes at the dimer interface are conserved across the IWA family^18^. The conformational landscape of other IWA ATPases such as the tail-anchored (TA) membrane protein targeting factor Get3 as well as the nitrogenase Fe-protein NifH have been extensively characterized, and their respective catalytic cycles involve conformational switching of the dimer between a *‘closed’* state that is catalytically competent for ATP hydrolysis and an *‘open’* state that is ATPase inactive^20–25^. The *open* conformation is reported for either the apo or post-hydrolysis MgADP-bound states of the ATPase cycle and is characterized by P-loops that are far apart^24,26^, whereas the *closed* state is reported for the pre- or mid-hydrolysis states where the two P-loops are adjacent^20,21^, mediating the position of the IWA lysines and thereby regulating nucleotide hydrolysis across the dimer interface. Two additional loop motifs, Switch I and II (Fig. S1A-B), are conserved across IWA ATPases and adopt discrete conformations depending on the nucleotide state. These nucleotide-dependent conformational transitions are critical for the function of IWA ATPases as ATP binding and hydrolysis is functionally coupled to the targeting of a substrate specific to each enzyme^18^.

Biochemical and spectroscopic characterization of ArsA suggests that the ATPase undergoes a series of conformational changes through its catalytic cycle in the presence of nucleotide and metalloid^27,28^. Four X-ray crystal structures have been reported for *E. coli* ArsA (*Ec*ArsA) in multiple nucleotide-bound states^14,29^. In each of these structures, MgADP is bound at the nucleotide-binding site on the N-domain, whereas MgADP (PDB:1F48), MgATP (PDB: 1II0), MgAMP•PNP (PDB: 1II9), or MgADP•AlF_3_ (PDB: 1IHU) are bound at the analogous site on the C-domain. Despite the distinct nucleotide states, only subtle conformational changes are seen between the various structures with a root mean square deviation (r.m.s.d.) of 0.38 Å over 536 residues (Fig. S1C), supporting that crystallization has likely restricted the range of motion in the protein. How ArsA undergoes nucleotide-dependent conformational transitions through its catalytic cycle to facilitate As^III^ efflux remains unclear.

The presence of metalloid substrate activates ArsA by stimulating ATP hydrolysis, which is the proposed rate-limiting step of the enzyme^27^. The underlying mechanism for this enhancement is poorly understood. Biochemical studies with Sb^III^ suggest that the metalloid substrate binds at a single high-affinity site in ArsA^30^. Of the three conserved cysteines, Cys172, located on a flexible loop between helices 7 and 8 of the N-domain of *Ec*ArsA, has been proposed to control the metalloid affinity of this site^30,31^. As the crystal structures are not consistent with these results, the mechanism of metalloid binding and ArsA activation requires further investigation.

In this article, we report single-particle cryogenic electron microscopy (cryo-EM) structures of ArsA from a thermotolerant and acidophilic bacteria, *Leptospirillum ferriphilum*, in nucleotide-bound *open* and *closed* conformations. In the MgADP-bound *open* state, the ArsA pseudodimer adopts the conformation of the previously reported ArsA crystal structures, all representing an ATPase inactive *open* state. MgATP alone is unable to stabilize a major conformational change from this *open* state. In the presence of MgATP and As^III^, a distinct *closed* conformation is adopted that is analogous to the *closed* conformations observed for other IWA ATPases. These structures provide views of the conformational transitions in the ArsA catalytic cycle. Combined with X-ray absorption spectroscopy (XAS) analysis and supporting biochemical studies, our cryo-EM structures suggest the molecular mechanism of As^III^-activated ATP hydrolysis in ArsA. Altogether, this work provides a structural and biochemical foundation for understanding the mechanism of As^III^ efflux by the ArsAB system.

## Results

### ArsA ATPase from *Leptospirillum ferriphilum* strain ML-04

Following a bioinformatic analysis of *ars* operons resembling the well-studied *E. coli* R773 plasmid *ars* operon^3^, we identified an ArsA homolog from *Leptospirillum ferriphilum* strain ML-04 (*Lf*ArsA), an acidophilic and moderately thermophilic bacterial species, where some strains are found in arsenic-rich acid mine drainage ecosystems (Fig. S2A)^32^. The ArsA gene has been shown to express under arsenic stress conditions in this strain, and the operon confers arsenic resistance in closely related *L. ferriphilum* strains^32–34^. *Lf*ArsA has 69% sequence identity to *Ec*ArsA, retaining all the characteristic features of ArsA (Fig. S1B). We cloned, overexpressed, and purified *Lf*ArsA with a C-terminal 6x-His tag. Purified *Lf*ArsA hydrolyzes ATP *in vitro* with a pseudo first-order rate constant of 0.5 ± 0.1 min^−1^, and saturating concentrations of As^III^ stimulate the ATPase activity ∼15-fold above the basal activity (Fig. S2B, *black* plot). To elucidate the conformational landscape of the ArsA catalytic cycle, we sought to determine nucleotide-bound *Lf*ArsA structures both in the presence and absence of its substrate As^III^ using cryo-EM.

### Structure of the *open* conformation of ArsA bound to MgADP

To determine the conformation of ArsA in an MgADP and As^III^-bound state, ArsA was incubated with 2 mM MgCl_2_, 2 mM ADP, and 5 mM sodium arsenite, and grids were prepared for single-particle analysis. From a dataset of 4,732 movies collected on a 300 kV Titan Krios Transmission Electron Microscope (TEM) and several rounds of 2D and 3D classification (Fig. S3), we obtained a 103,258-particle reconstruction of the ArsA•MgADP state (Fig. 1A). The final Coulomb potential map was refined to a 3.4 Å gold-standard Fourier shell correlation (FSC) resolution and sharpened using a uniform B-factor of −165 Å^2^ to better visualize high-resolution features. AlphaFold2^35^ predictions for the N- and C-domains of *Lf*ArsA were docked into the sharpened map, followed by model building and refinement to obtain the ArsA•MgADP structure (Fig. 1B). The average Q-score for the structure was 0.64 (Fig. S4A), indicating good resolvability of the cryo-EM map^36^. About 95% of the protein chain was modeled into the map, with missing residues including part of the linker region between the N- and C-domains (residues 295-307), residues 167-168 and residues 464-476 (Fig. 1B). The N (residues 1-298) and C (residues 315-587) domains are pseudo-symmetric with an r.m.s.d. of 1.0 Å over 139 residues (Fig. 1C). At both nucleotide-binding sites, ADP is coordinated to a Mg^2+^ ion (Fig. 1D). The catalytic Switch I aspartates – Asp46 (N) and Asp364 (C) – are oriented away from the Mg^2+^ ion (>4.5 Å). The Switch II aspartates – Asp143 (N) and Asp447 (C) – interact with the Mg^2+^ ion within 4 Å and form short hydrogen bonds with Thr23 (N) and Thr341 (C), respectively. Notably, the P-loops are sufficiently separated such that the catalytic IWA lysines (Lys17 and Lys335) are oriented away from the ADP bound at the opposing domain, positioning the terminal amino groups of the lysines over 8 Å away from the nucleotide phosphate groups (Fig. 1E).

**Figure 1.**
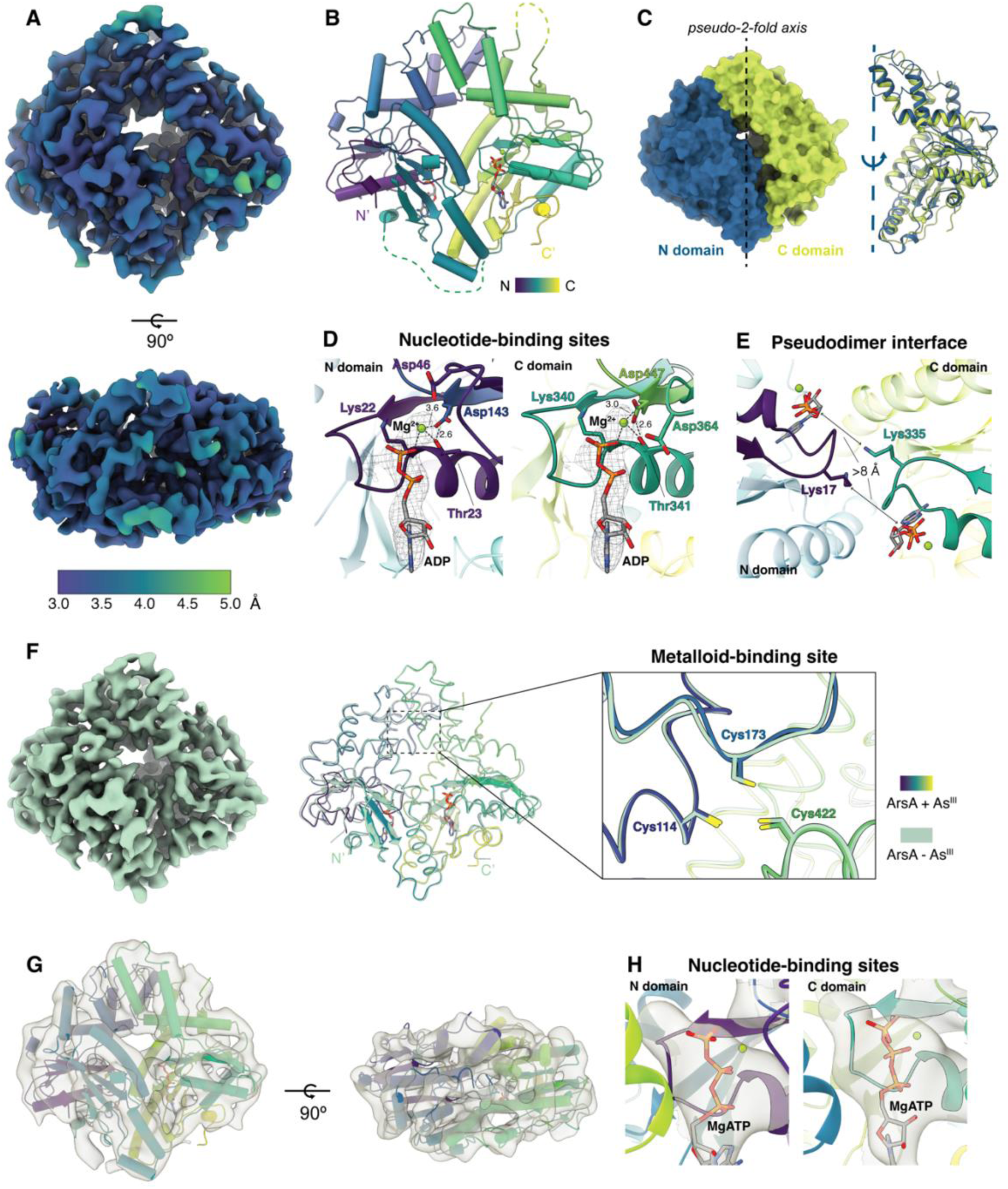
Cryo-EM structures of *Lf*ArsA in the nucleotide-bound *open* conformation. **(A)** Two views of the sharpened cryo-EM map colored based on local resolution (3.4 Å overall) for *Lf*ArsA•MgADP *open* state (*ADP-open*) in the presence of As^III^. **(B)** Cartoon representation of the *ADP-open* state colored in viridis. Unmodeled regions are indicated as dotted lines. **(C)** Left, surface representation of the ArsA pseudodimer highlighting the N-(blue) and C-(green) domains. Right, cartoon representation of the overlay of the two domains based on the pseudo - 2-fold symmetry. **(D)** Nucleotide-binding sites of the *ADP-open* state showing the Coulomb potential map around the MgADP shown as sticks. Switch I & II Asp residues of both N- and C-domains are shown as sticks. **(E)** The pseudo-dimer interface of the *ADP-open* state highlighting the distance between the IWA lysines. **(F)** Structure of *Lf*ArsA•MgADP *open* state in the absence of As^III^. Left, 3.8 Å cryo-EM map; middle, ribbons model (pale green) overlayed with the *ADP-open* state model from *B*; right, the metalloid-binding sites from both structures. **(G)** Cryo-EM map (6-7 Å overall resolution) of the *Lf*ArsA non-hydrolyzing variant (D46N/D364N) solved in the presence of MgATP with the *ADP-open* state model from *B* docked in the map. **(H)** Nucleotide-binding sites of the *Lf*ArsA•MgATP state showing Coulomb potential map likely corresponding to ATP. MgATP was manually fit using Coot.

We propose that our ArsA•MgADP structure represents an *open* form of ArsA, designated as the *‘ADP-open’* state for the rest of the discussion. Based on the general mechanism of the IWA ATPase family, this conformation represents an inactive form of ArsA unable to hydrolyze ATP. Comparison of the cryo-EM structure to the X-ray crystal structures of *Ec*ArsA reveals the structures adopt similar conformations (Fig. S1C); for example, the cryo-EM structure and the *Ec*ArsA•MgADP crystal structure (PDB: 1F48) exhibit an r.m.s.d. of 0.95 Å over 475 residues. Despite the presence of millimolar concentrations As^III^ in the samples, there was no evidence of As^III^ at the metalloid-binding site in either our EM or the X-ray structures (Fig. S5)^14,29^.

### Structures of nucleotide-bound ArsA in the absence of As^III^

To clarify the absence of As^III^ at the metalloid-binding site in the ArsA•MgADP structure, we solved a cryo-EM structure using identical experimental conditions but in the absence of As^III^. The resulting 3.8 Å resolution map of the ArsA•MgADP state is in *open* conformation (Fig. 1F, S4B & S6), similar to the structure solved in the presence of As^III^, with an r.m.s.d. of 0.57 over 545 residues. Importantly, the configuration of the metalloid-binding site remains unchanged (Fig. 1F; inset on right), confirming that both states lack As^III^. This suggests that the *ADP-open* state does not effectively bind the metalloid.

The presence of As^III^ stimulates steady-state ATP hydrolysis by ArsA above the basal activity (Fig. S2B)^28,30^. To characterize the ATP-bound ArsA conformation in the absence of metalloid, we generated a non-hydrolyzing variant of *Lf*ArsA by mutating the catalytic Switch I aspartates – Asp46 and Asp364 of the N- and C-domains, respectively – to asparagines (Fig. S2B, *red* plot). We used this variant to capture the MgATP-bound state, similar to the approach used to obtain the MgATP-bound Get3 structure (PDB: 7SPY)^21^. For the rest of the discussion, structures solved using the D46N/D364N variant will be simply referred to as ArsA. Under experimental conditions similar to those used to solve the *ADP-open* structures, we could only obtain a low-resolution (6-7 Å) EM map that revealed an *open* state of ArsA (Fig. S7). We could confidently fit the model of the ArsA•MgADP structure into the map (Fig. 1G). While the density observed at the nucleotide-binding sites could not be unambiguously distinguished at this resolution, it is consistent with MgATP binding (Fig. 1H). This is reminiscent of NifH that is refractory to crystallization in the presence of ATP but readily crystallizes in the presence of both ATP and its partner protein, NifDK (MoFe protein)^26^. It is plausible that binding of ATP makes ArsA conformationally flexible, preventing high-resolution cryo-EM reconstruction, and that a partner protein or As^III^ is required for stabilization. Importantly, this structure is consistent with the low basal ATPase activity of ArsA. Together, these As^III^-free structures support that nucleotide binding alone is unable to stabilize a conformation competent for ATP hydrolysis.

### Structure of the *closed* conformation of ArsA bound to MgATP and As^III^

To understand the structural basis of As^III^ activation, we sought to characterize the conformation of ArsA in the presence of both MgATP and As^III^. We incubated the non-hydrolyzing ArsA variant with 2 mM MgCl_2_, 2 mM ATP, and 2 mM sodium arsenite and then performed single-particle cryo-EM (Fig. S8). The resulting Coulomb potential map was resolved to an overall gold-standard FSC resolution of 3.0 Å and sharpened using a uniform B-factor of −124 Å^2^ (Fig. 2A). AlphaFold2 models of N- and C-domains of *Lf*ArsA were docked into the sharpened density map, the model was built and refined to obtain the structure of ArsA•MgATP•As^III^ with clear density for MgATP at both nucleotide-binding sites (Fig 2B-C). Significantly, at the metalloid-binding site, we observed density enclosed between the three conserved cysteines – Cys114, Cys173, and Cys422 – into which we modeled an arsenic atom (Fig. 2D). The average Q-score for the structure was 0.71 (Fig. S4C), indicating high resolvability of the cryo-EM map. About 97% of the protein was modeled in, while the regions with poor density, such as most of the linker region (residues 299-305) and residues 474-479, were omitted from the final model.

**Figure 2.**
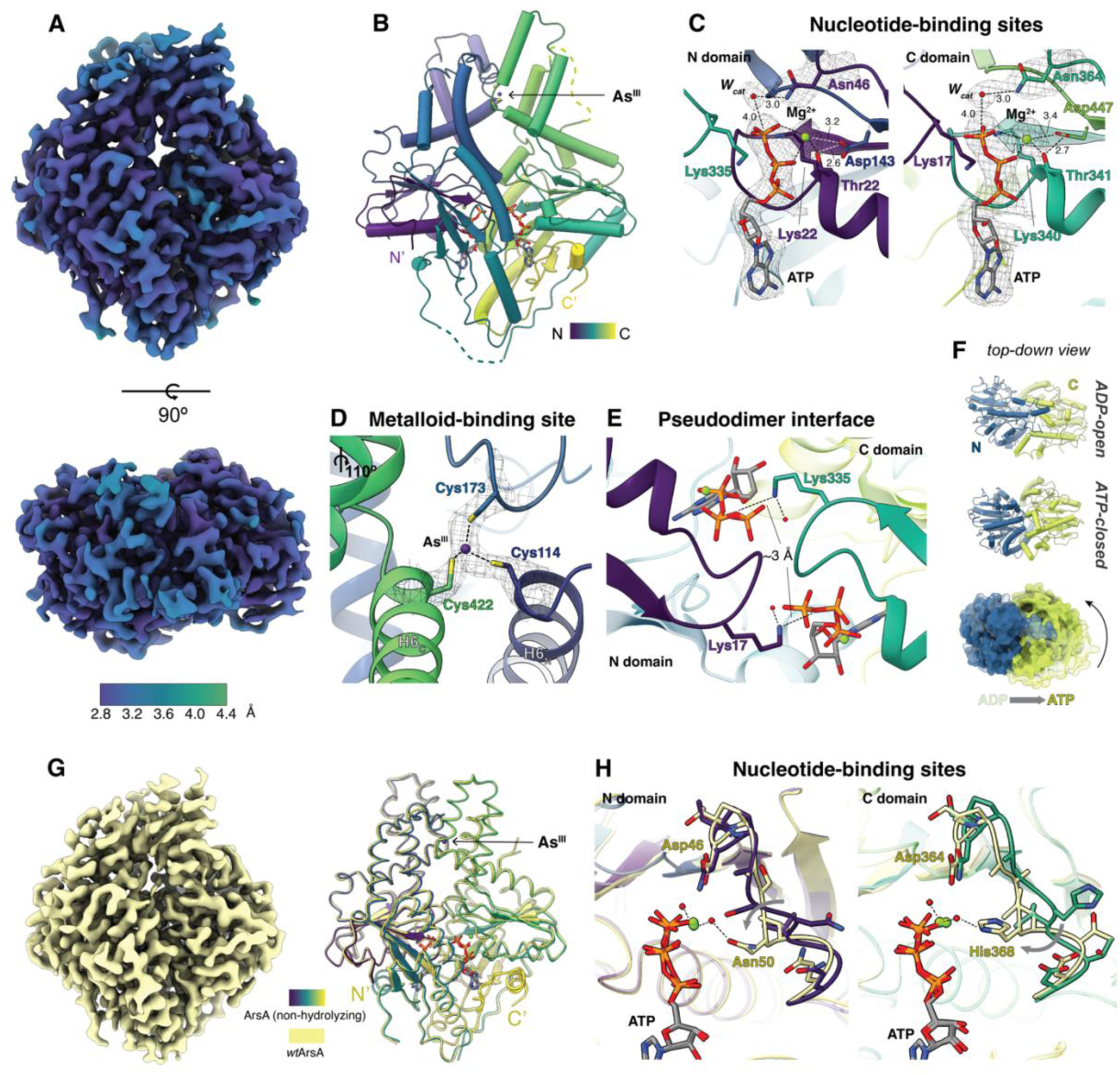
Cryo-EM structure of *Lf*ArsA *closed* conformation bound to MgATP and As^III^. **(A)** Two views of the sharpened cryo-EM map colored based on local resolution (3.0 Å overall) for *Lf*ArsA•MgATP•As^III^ *closed* state (*ATP-closed*) of the non-hydrolyzing variant (D46N/D364N). **(B)** Cartoon representation of the *ATP-closed* state colored in viridis. Unmodeled regions are indicated as dotted lines. **(C)** Nucleotide-binding sites of the *ATP-closed* state showing the Coulomb potential map around MgATP and Switch I catalytic residues (Asn46 and Asn364) shown as sticks. Switch II Asp residues of both N- and C-sites are shown as sticks. The nucleophilic water molecule (*W_cat_*) is highlighted. **(D)** Metalloid-binding site of *ATP-closed* state showing Coulomb potential map for As^III^ (purple atom) coordinated by Cys114, Cys173 and Cys422. The view is rotated 110° clockwise relative to *B*. **(E)** The pseudodimer interface of the *ATP-closed* state highlighting the distance between the IWA lysines (Lys17 and Lys335). **(F)** Top-down view, relative to *A*, comparing the *ADP-open* and the *ATP-closed* states of *Lf*ArsA colored as in Fig. 1C. showing both states in cartoon and then overlayed in surface representation. The overlay is aligned to the N-domain P-loop (residues 16-23) showing the relative change of the C-domain. **(G)** Structure of *Lf*ArsA•MgATP•As^III^ *closed* state solved using wild-type *Lf*ArsA. Left, 3.0 Å map cryo-EM map; right, ribbons model (yellow) overlayed with the model from *B*. **(H)** Comparison of Switch I conformations between the two *ATP-closed* structures. Differences in interactions are highlighted.

Within both nucleotide-binding sites, ATP is stabilized by interactions of its γ-phosphate group with Mg^2+^ and Lys (22 and 340, respectively) by electrostatic interactions and the ⍺- and β-phosphates with the P-loop backbone by hydrogen bonds (Fig. 2C & Fig. S9A). The ribose sugar and the adenine base are stabilized by hydrogen bonds contributed by sidechains from both domains and backbone hydrogen bonds from the loop region corresponding to the adenine-binding loop (A-loop) described in Get3 structures (Fig. S9A)^24^. Additionally, ATP bound to the N-domain is stabilized by potential cation-π interactions formed between the adenine base and both Arg207 (N) and Arg544 (C) (Fig. S9A, left). However, in the C-domain, the corresponding interaction is only observed between the adenine and Arg256 (N). In this structure, the Switch I aspartates (mutated to asparagines here) – Asn46 (N) and Asn364 (C) – are positioned adjacent to each Mg^2+^ ion and anchor the catalytic water molecule (*W_cat_*) near the *γ*-phosphate of ATP via a 3.0 Å hydrogen bond for nucleophilic attack, priming the enzyme for hydrolysis (Fig. 2C). Switch II aspartates – Asp143 (N) and Asp447 (C) – are positioned at 3.2 and 3.4 Å from the Mg^2+^ ion respectively and form short hydrogen bonds with the Thr residue coordinated to the respective Mg^2+^ ions, similar to the MgADP-bound structure.

This structure adopts a *closed* pseudodimer conformation previously unobserved for ArsA that we define as the *‘ATP-closed’* state. Notably, the P-loops of the two domains are in close proximity (Fig. 2E). The terminal amino group of the conserved IWA lysine of each domain – Lys17 (N) and Lys335 (C) – is positioned ∼3 Å from the bridging oxygen atom between the β- and γ-phosphate groups of the ATP bound to the opposing domain. This enables stabilization of negative charge build-up on the nucleotides in the transition state. Together with the Switch I aspartates, which are positioned to activate *W_cat_* for nucleophilic attack on ATP, this *closed* pseudodimer conformation represents a catalytically competent state of ArsA at both nucleotide-binding sites. The *ATP-closed* state is analogous to the *closed* dimer states previously reported for Get3 (PDB: 7SPY) and NifH (PDB: 1M34) (Fig. S10)^21,37^. Notably, in our structure, similar to the Get3 *closed* structure, both IWA lysines appear to form a hydrogen bond with a water molecule found at the pseudodimer interface; these waters may assist with anchoring the terminal amino groups close to the nucleotides in preparation for ATP hydrolysis (Fig. 2E).

Relative to the *ADP-open* state, the two domains rotate towards the pseudodimer interface, forming a tighter interface in the *ATP-closed* state (Fig. 2F). This generates new inter-domain contacts not observed in the *ADP-open* state (Fig. S8B). Glu215 and Glu507, from the N- and C-domains, respectively, form hydrogen bonds with the catalytic water molecule and Switch I residues of the opposing nucleotide-binding site. Electrostatic contacts are formed between Arg214 of the N-domain and Asp371 of the C-domain. This results in a buried surface area of 2,380 Å^2^, ∼500 Å^2^ more than the *open* conformation.

We next sought to establish the conformation of ArsA under turnover conditions. Wild-type ArsA (*wt*ArsA) was incubated with 2 mM MgCl_2_, 2 mM ATP, and 2 mM sodium arsenite at room temperature for ∼1.5 min before cryo-EM grid preparation. This resulted in a Coulomb potential map at 3.0 Å overall gold-standard FSC resolution (Fig. 2G, left; S4D & S11). The final refined structure revealed a *closed* conformation similar to the ArsA non-hydrolyzing variant (r.m.s.d = 0.53 Å over 565 residues) with MgATP present at both nucleotide-binding sites and arsenic modeled at the metalloid-binding site (Fig. 2G, right). The IWA lysines in this structure are oriented towards the nucleotides (Fig. S12A), supporting a catalytically competent pre-hydrolysis state consistent with the variant structure. While the Switch I aspartate in each nucleotide-binding site is positioned next to the Mg^2+^ ion anchoring the *W_cat_* for nucleophilic attack, subtle differences are observed in the conformations of the Switch I loop between the *wt*ArsA and variant *closed* structures. In the N-domain, the Switch I loop moves towards the nucleotide in *wt*ArsA such that Asn50 now interacts with the Mg^2+^ ion via an intervening water molecule (Fig. 2H, left). Likewise, in the C-domain, Switch I loop shifts towards the nucleotide, causing His368 to interact with Mg^2+^ via an intervening water molecule (Fig. 2H, right). Additionally, we see evidence for a hydrogen-bonding network composed of ordered water molecules including *W_cat_*, that connects the Switch I aspartates at the N-(Asp46) and C-(Asp364) sites, potentially coupling the two active sites (Fig. S12B). Together, these subtle structural changes observed in the pre-hydrolysis state prime the enzyme to initiate hydrolysis in both N- and C-domains.

### The As^III^-binding site of ArsA

We observed density for arsenic at the metalloid-binding site in the *ATP-closed* state, coordinated by the thiolate groups of Cys114, Cys173, and Cys422 in a pyramidal geometry (Fig. 2D and S12C). Cys114 and Cys422 are homologous residues within the pseudodimer, located at the N-terminal end of helix 6, denoted as H6_N_ and H6_C_, respectively, while Cys173 is located on a flexible loop (residues 155-183) between helix 7 and helix 8 in the N-domain.

To confirm the As^III^ coordination environment, we performed solution X-ray absorption spectroscopy (XAS). ArsA non-hydrolyzing variant (1.95 mM) supplemented with 15 mM MgCl_2_, 15 mM ATP, and 1.5 mM sodium arsenite was subjected to XAS analysis. X-ray absorption near edge spectra (XANES) confirmed the presence of trivalent arsenic (Fig. 3A, *black* spectra), as the K-edge at 11865.9 eV was consistent with the first inflection edge energies of As^III^ model compounds (11867 eV). Best-fit simulations of the extended X-ray absorption fine structure (EXAFS) spectra supported the presence of As^III^ coordinated by three sulfur ligands (AsS_3_) at an average As-S bond length of 2.27 Å (Fig. 3B, top panel; Table S2), consistent with cysteine coordination in the cryo-EM structure. Long-range carbon scattering was also observed in the sample, likely from secondary and tertiary sphere carbon atoms associated with the β- and ⍺-carbon atoms of the cysteines. The oxidation state and coordination environment of the ArsA sample are consistent with that of a control sample consisting of As^III^ mixed with L-cysteine (Fig. 3A, *purple* spectra & Fig. 3B, middle panel; Table S2). Similar As^III^ coordination has been reported for the As^III^-responsive DNA repressor, ArsR, and the As^III^ metallochaperone, ArsD, based on XAS analysis^38–40^. We quantified the amount of As^III^ bound to ArsA under conditions resembling those used in cryo-EM sample preparation by inductively coupled plasma mass spectrometry (ICP-MS). In the presence of MgATP, ArsA binds As^III^ with 1:1 stoichiometry (Fig. 3C, *black* plot). Taken together, cryo-EM, XAS, and ICP-MS analysis support the binding of one three-coordinate As^III^ at the ArsA pseudodimer interface in the presence of MgATP.

**Figure 3.**
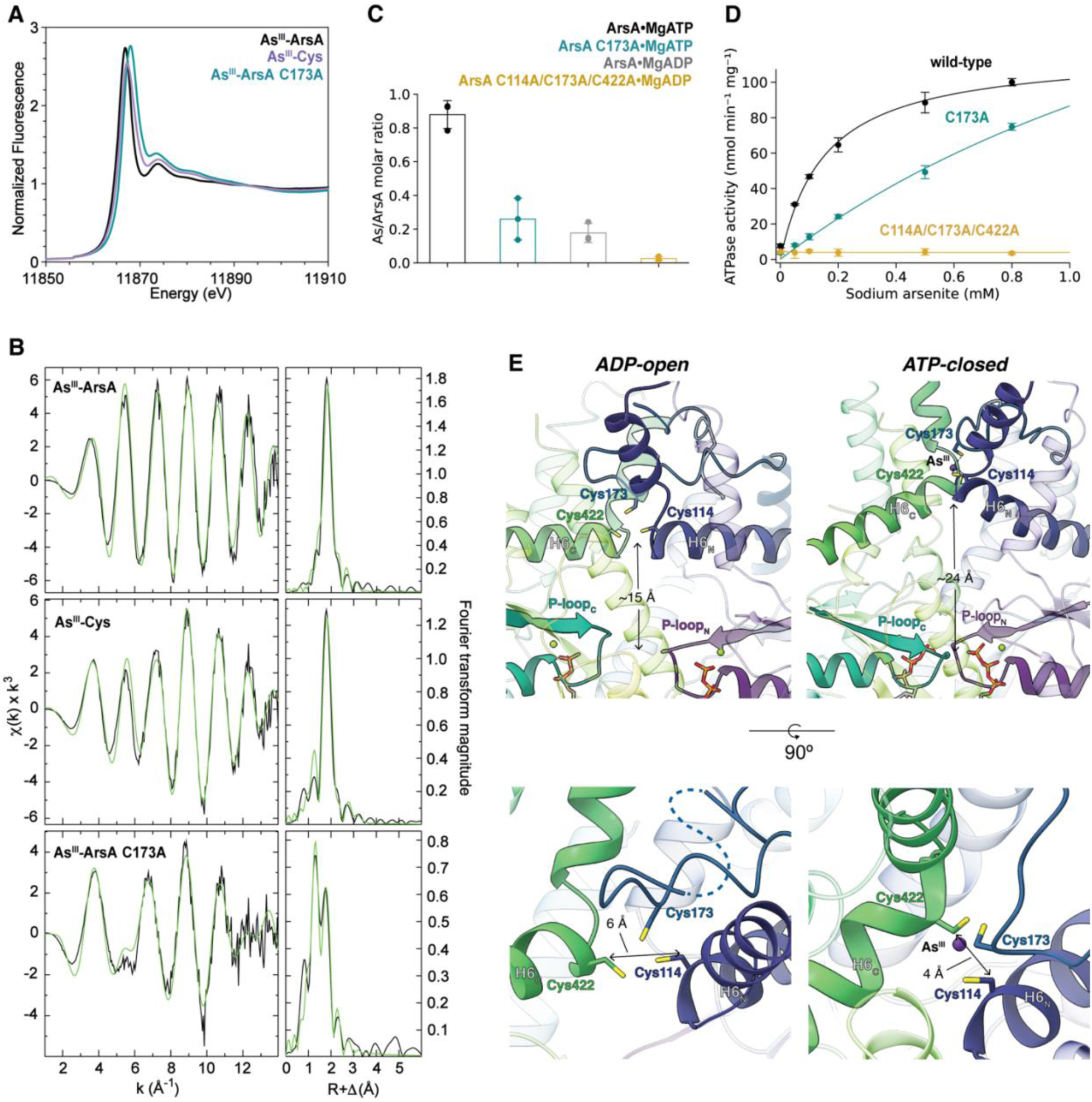
Characterization of the As^III^ binding site of ArsA. **(A)** X-ray absorption near edge spectra (XANES) for *Lf*ArsA•MgATP•As^III^ (‘As^III^-ArsA’; black), As^III^-L-cysteine control (‘As^III^-Cys’; purple), and *Lf*ArsA(C173A)•MgATP•As^III^ (‘As^III^-ArsA C173A’; teal). **(B)** Left column, extended X-ray absorption fine structure (EXAFS) spectra (black) with best-fit simulations (green) for each complex. Right column, corresponding Fourier-transforms. A replicate of each XAS sample was individually prepared and analyzed (Fig. S13). **(C)** ICP-MS analysis for As^III^ quantification in nucleotide-bound *Lf*ArsA samples (ArsA•MgATP, blac*k*; ArsA C173A•MgATP, teal; ArsA•MgADP, gray; ArsA C114A/C173A/C422A•MgADP, yellow). Bars represent mean As/ArsA molar ratio for *n* = 3 individually prepared samples with error bars representing standard deviation. **(D)** Steady-state ATPase activity of *Lf*ArsA (nmol min^−1^ mg^−1^) plotted against varying As^III^ concentrations (mM) in the presence of 5 mM MgCl_2_ and 5 mM ATP at 37°C (wild-type, black; C173A, teal; C114A/C173A/C422A, yellow). Data points represent mean of n = 3 and error bars represent standard deviation. The data was fit to the Michaelis-Menten equation. **(E)** Comparison of metalloid-binding sites of *ADP-open* (left) and *ATP-closed* (right) highlighting the relative positions of Cys114, Cys173 and Cys422 that enable As^III^ (purpl*e*) binding.

The As^III^-coordinating cysteines are conserved among ArsA homologs (Fig. S14). Alanine mutations of these residues ablate As^III^ activation of steady-state ATP hydrolysis (Fig. 3D*, yellow* plot). By contrast, mutation of Cys173 located on the flexible loop region of N-domain alone appears to only decrease the binding of As^III^ to ArsA, as the C173A variant requires higher As^III^ concentrations than the wild-type enzyme to approach half-maximal activation of ATPase activity (Fig. 3D, *teal* plot). XANES and EXAFS spectra for the C173A variant incubated with MgATP and As^III^ suggest AsO_2_S coordination, indicating that As^III^ is no longer coordinated to three cysteines (Fig. 3A, *teal* spectra & Fig. 3B, bottom panel; Table S2). Moreover, ICP-MS analysis of this variant shows over 50% decrease in As^III^-ArsA molar ratio (Fig. 3C, *teal* plot). As previously proposed^30^, Cys173 regulates the binding affinity of As^III^ and is not critical for the stimulation of nucleotide hydrolysis.

Based on our ICP-MS analysis, ArsA•MgADP binds As^III^ far less tightly than ArsA•MgATP (Fig. 3C, *black* and *gray* plots), corroborating the absence of As^III^ density in the *ADP-open* structure. Mutating the As^III^-coordinating cysteines completely disrupts As^III^ binding to ArsA•MgADP (Fig. 3C, *yellow* plot). How do the conformational changes between *open* and *closed* states modulate the affinity for As^III^? Our cryo-EM structures reveal that switching from *open* to *closed* state causes reorientation of helix 6 of both domains such that Cys114 on H6_N_ and Cys422 on H6_C_ shift ∼9 Å away from the P-loops (Fig. 3E, top panels). This makes the metalloid-binding site more accessible to the solvent for As^III^ binding. Additionally, a change in the relative positions of the sidechains of Cys114 and Cys442 accompanies this conformational rearrangement. As shown in Fig. 3E (bottom panels), the distance between C_β_ atoms of Cys114 and Cys422 reduces from 6 Å in the *open* state to 4 Å in the *closed* state, facilitating As^III^ binding. Therefore, by controlling the relative positions of the three cysteines at the metalloid-binding site, conformational changes in ArsA modulate the binding affinity for As^III^.

### Intra-domain conformational rearrangements between *open* and *closed* states

In addition to the nucleotide-dependent conformational changes at the pseudodimer interface, large-scale structural rearrangements are observed within each domain between the *ADP-open* and the *ATP-closed* states (Fig. 4A-B). These changes are primarily linked to the Switch I and II motifs. Compared to the *ADP-open* state, the Switch I loop of both the N- and C-domains rearranges to shift towards the nucleotide in the *ATP-closed* state (Fig. 4C-D). This enables the positioning of Asp46 and Asp364 to activate *W_cat_* and initiate ATP hydrolysis. Differences in the Switch I conformation between the two domains allow Ser49 to additionally stabilize the catalytic water in the N-domain (Fig. 4C & G). While the carboxylate group of Asp364 shifts about 5 Å towards the active site from *open* to *closed* state, the carboxylate of Asp46 shifts only 3 Å (Fig. 4C-D).

**Figure 4.**
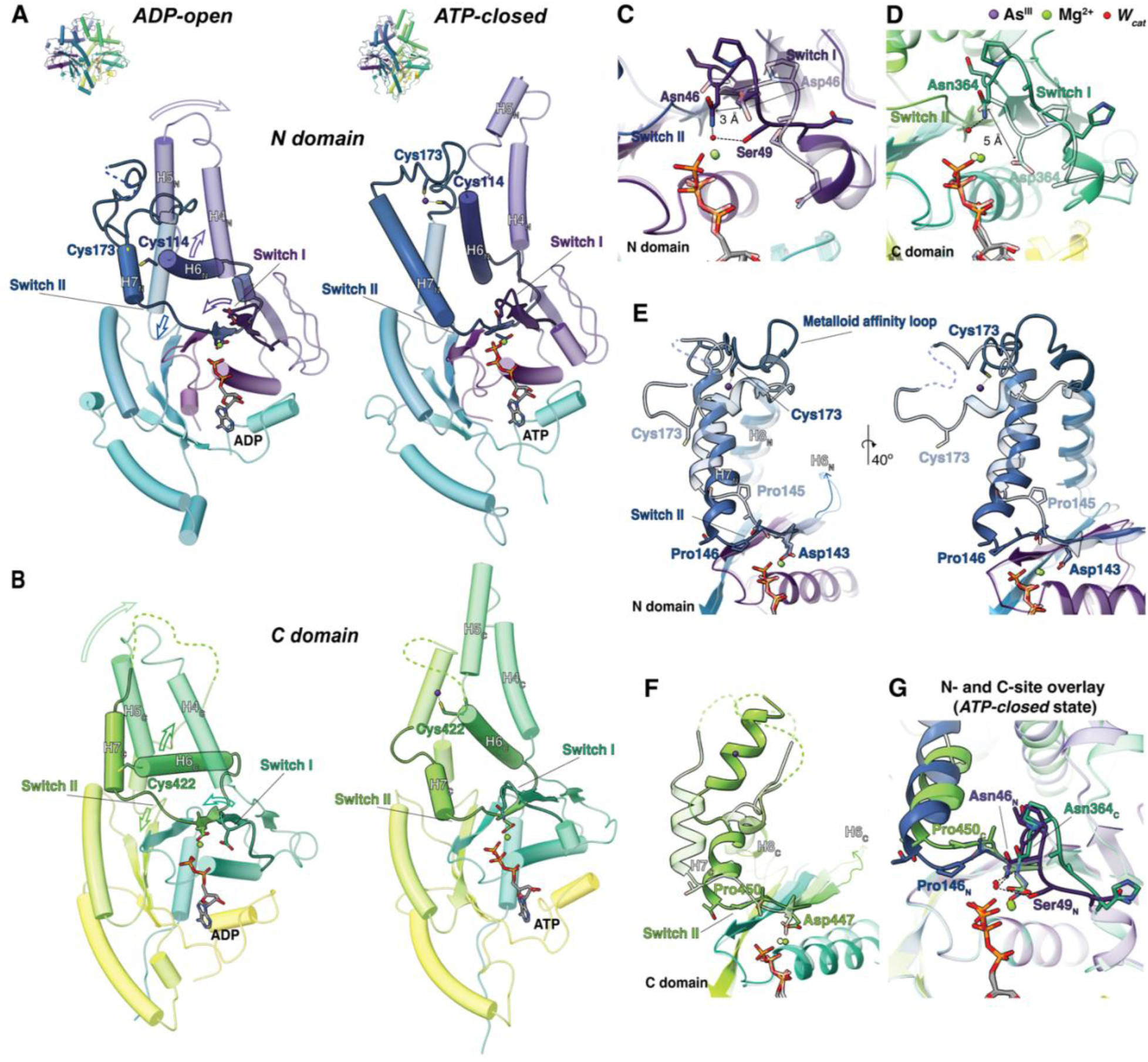
Intra-domain conformational changes between *ADP-open* and *ATP-closed* states. **(A,B)** Cartoon representations of overall structural rearrangements within N- and C-domains. Arrows indicate direction of motion of the corresponding motif from the *open* to the *closed* state. This and subsequent panels are colored in viridis. Highlighted residues are shown as sticks. All overlayed structures are aligned to the respective P-loops (residues 16-23 (N) and 334-341 (C)) and the transparent structures represent the *ADP-open* state. **(C, D)** Conformational changes in Switch I. **(E)** Conformational changes in Switch II and H7_N_ of the N-domain. The ‘metalloid affinity loop’ connecting H7_N_ and H8_N_ is highlighted. In both panels E and F, helix 6 has been omitted for visual clarity. **(F)** Conformational changes in Switch II and H7_C_ of the C-domain. **(G)** Comparison of Switch I and II conformations of the N- and C-domains in the *ATP-closed* state.

Switch II, acting as a link between the nucleotide-binding site and the metalloid-binding site, modulates a series of dramatic conformational changes within each domain between the *open* and *closed* states. Upon As^III^ binding to the metalloid-binding site of the *ATP-closed* state, both H6_N_ and H6_C_ undergo rigid body rearrangement such that the N-terminal end of each, bearing the As^III^ ligands – Cys114 and Cys422, respectively – rotates away from the nucleotide-binding sites (Fig. 4A-B). Reorientation of helix 6 leads to restructuring of the downstream Switch II motif in the *ATP-closed* state. Although the Switch II aspartates (Asp143 and Asp447) of the respective domains remain anchored near the Mg^2+^ ion in both states, downstream residues 144-147 of the N-domain (Switch II_N_), residues 448-451 of the C-domain (Switch II_C_), and both helix 7s shift towards the nucleotide in the *ATP-closed* state (Fig. 4E-F). Switch II_N_ shifts towards the nucleotide to a greater extent than Switch II_C_ (Fig. 4G). Hydrogen-bonding interactions between Thr147 (Switch II_N_) and Glu117 (H6_N_) and between Thr451 (Switch II_C_) and Glu425 (H6_C_), in the *ADP-open* state, are destabilized in the *ATP-closed* state to facilitate the conformational change coordinated by helix 6 and Switch II (Fig. S15). In the *wt*ArsA *ATP-closed* state, Switch II_C_ further shifts towards the nucleotide, adopting the same conformation as Switch II_N_ (Fig. S12D). This symmetric conformation of Switch II results in an additional contact between the two domains where His149 (H7_N_) and His453 (H7_C_) form a π-π stacking interaction (Fig. S12E).

A striking difference is seen between the N- and C-domains in the conformation of helix 7 following the Switch II loop. In the *ADP-open* state, H7_N_ is composed of residues Gly148-Leu154. In the *ATP-closed* state, this helix is extended by incorporation of formerly loop residues Gln155-Ala166 into the C-terminal end of H7_N_ transition, resulting in a twisted helix composed of residues Thr147-Ala166 (Fig. 4E). By contrast, H7_C_ transitions from residues Gly452-Ala460 in the *ADP-open* state to a discontinuous helical segment between Thr451 and Gln471, connected by a loop formed by residues Asp459-Gly462 in the *ATP-closed* state (Fig. 4F). Complete helical rearrangement of H7_N_ arises by virtue of Cys173 on the loop region between H7_N_ and H8_N_, that serves as the third ligand for As^III^ binding in the *ATP-closed* state. This loop, designated as the ‘metalloid affinity loop’, undergoes a considerable rearrangement between the *open* and *closed* states to stabilize three-coordinate As^III^ binding (Fig. 4E). An analogous residue is not found on the C-domain; in fact, the corresponding loop region in the C-domain is poorly resolved in both states. Notably, such helical transition of helix 7 is also observed upon substrate (TA protein) binding at the homologous site in Get3 (PDB: 7SQ0)^21^. Taken together, the conformational changes associated with As^III^ binding in the *ATP-closed* state demonstrate how three-coordinate As^III^ binding at the metalloid-binding site of ArsA is allosterically coupled to the nucleotide-binding sites via Switch II.

## Discussion

ArsA enhances the efficiency of ArsB by coupling ATP hydrolysis to toxic metalloid (As^III^ or Sb^III^) efflux. Based on pre-steady state kinetic analysis, the pseudodimeric ATPase undergoes nucleotide-dependent conformational changes throughout its catalytic cycle^27,28,41^, that are critical for its interaction with ArsB. MgATP-bound ArsA has been reported to alternate between two conformations that differ in their affinities for the metalloid substrate^27^. Metalloid binding stabilizes one of these conformations, followed by hydrolysis and phosphate release. Our high-resolution cryo-EM efforts provide the structural basis for the conformational states proposed in this mechanism. We have characterized *open* and *closed* conformations of ArsA. The *open* state can exist bound to either ADP or ATP (Fig 1B, F-G), has low metalloid affinity, and is ATPase inactive. The *closed* state is stabilized as a ternary complex when As^III^ binds ArsA•MgATP (Fig. 2B) and is competent for ATP hydrolysis. The structures enable us to propose a catalytic mechanism for ArsA (Fig. 5). ATP binds both N- and C-sites of ArsA where the enzyme exists in equilibrium between *open* and *closed* states (states 1 and 2). ATP alone is unable to drive the equilibrium to favor the *closed* state; as a result, ArsA hydrolyzes nucleotide at a basal rate. When As^III^ is coordinated by three cysteines at the metalloid-binding site (Fig. 2D), the *open* conformation transitions into the catalytically competent *closed* conformation (state 3). Coordination of As^III^ to Cys114 and Cys422 allosterically triggers the transition of the pseudodimer from *open* to *closed* state via helix 6 and Switch II (Fig. 4A-B). Cys173 on the ‘metalloid affinity loop’ acts as a switch controlling the binding affinity of As^III^ for ArsA and stabilizes the ternary complex in the *closed* state. As^III^ binding by this motif is coupled to the nucleotide-binding site of the N-domain via H7_N_ and Switch II_N_ (Fig. 4E). This mechanism reveals how three-coordinate As^III^ binding allosterically stimulates ATP hydrolysis, which is the proposed rate-limiting step of ArsA^27^. Following ATP hydrolysis, the enzyme loses affinity for As^III^, releasing the metalloid and phosphate, and returns to the *open* state (state 4). Nucleotide exchange can then reset the enzyme for another catalytic cycle.

**Figure 5.**
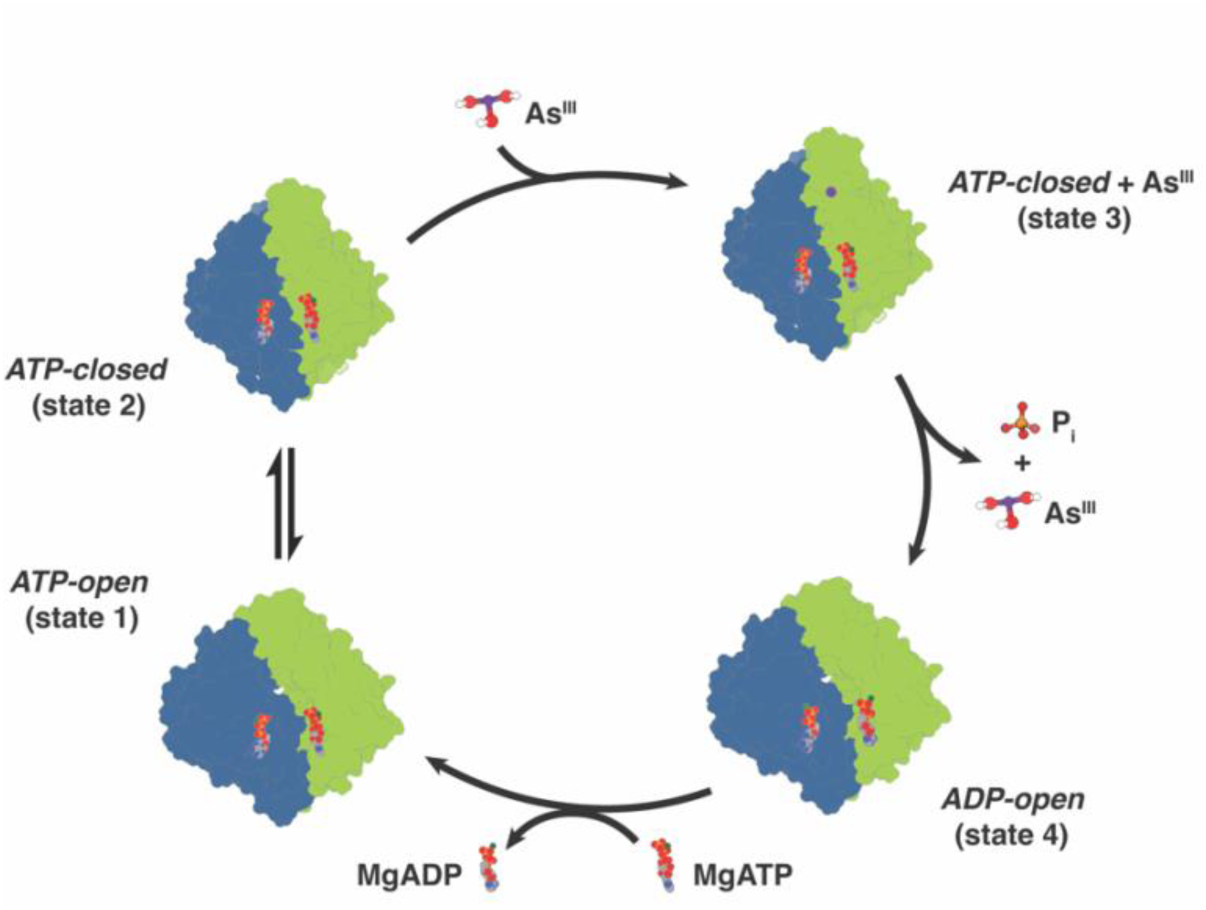
Mechanistic model for ArsA catalytic cycle. ATP-bound ArsA alternates between *open* (state 1) and *closed* (state 2). As^III^ binding stabilizes the catalytically competent *closed* conformation (state 3). Following ATP hydrolysis and release of As^III^ and phosphate (P_i_), ArsA switches to the ADP-bound *open* conformation (state 4). Exchange of the nucleotide resets the enzyme for another catalytic cycle. States 1, 3 and 4 were characterized in this study. ArsA is colored by N (blue) and C (green) domains. Nucleotides are shown as space-filling models. As^III^ (As(OH)_3_ at physiological pH) and P_i_ are shown as ball & sticks.

The conformational changes of ArsA reported here are consistent with the general structural and mechanistic features of IWA ATPases. Despite each IWA ATPase binding a unique substrate at the (pseudo)dimer interface, the structural changes corresponding to the substrate, *e.g.,* As^III^ binding to ArsA, remain conserved. In the case of Get3, TA protein binding induces a conformational change in helix 6 (substrate helix) and Switch II, thus regulating the dimer dynamics^21,25^. Likewise in NifH, association with NifDK affects the positioning of the [4Fe:4S] cluster for electron transport, which then induces conformational changes in the substrate helix and Switch II, stabilizing the *closed* dimer conformation^20^. Notably, the pseudodimeric architecture of ArsA is distinct from other ATPases in the family. While both N- and C-domains have been shown to be competent for ATP hydrolysis^17^, structural differences between the domains in the *ATP-closed* structures highlight the functional asymmetry within ArsA. Both Switch I and II loops adopt distinct conformations relative to the nucleotide in the N- and C-domains in the *closed* state (Fig. 4G & S16A), reflecting the difference in ATPase activities, where the N-site is more competent for hydrolysis than the C-site (Fig. S16B). These structural differences become less obvious in the *wt*ArsA *closed* state under turnover conditions where a potential hydrogen-bonding network linking the two active sites may coordinate hydrolysis at the two sites (Fig. S12B & D).

A key gap in our understanding of the ArsAB efflux pump is how the conformational landscape of the ArsA catalytic cycle modulates interactions with ArsB to facilitate As^III^ efflux. ArsB is sufficient to transport the metalloid and coupling the process with ATP hydrolysis enhances the efficiency of efflux^7^. A mutation in the N-domain P-loop of *Ec*ArsA (G20S) has been shown to abolish ATPase activity and inhibit the ability of ArsB to remove As^III^ from the cell^42^. This suggests that ArsA catalytic cycle is required for interaction with ArsB. In this light, nucleotide-dependent conformational changes between the high As^III^-affinity *closed* state and low As^III^-affinity *open* state uncovered in this study ensure efficient metalloid transfer to ArsB. This mechanism enables the survival of the bacteria under high arsenic stress conditions.

## Methods

The *Lf*ArsA gene was designed and purchased from Twist Biosciences (San Francisco, CA). Primers for amplification were designed using the New England Biolabs (NEB) (Ipswich, MA) primer design tool and obtained from Integrated DNA Technologies (Newark, NJ). Cloned constructs were sequenced by whole plasmid sequencing through Plasmidsaurus (Eugene, OR).

### Cloning, Expression, and Purification of LfArsA and its variants

*Lf*ArsA (Uniprot Accession Number: J9ZFA3) was cloned into the multiple cloning site-1 of pETDuet-1 expression vector encoding a 6x-His tag on the 3’-end, using Hi-Fi DNA assembly protocol (NEB). This resulted in an *Lf*ArsA construct with a C-terminal 6x-His tag. The vector backbone and gene fragment were amplified using the Q5 High-Fidelity 2X Master Mix protocol (NEB). ArsA variants were prepared by point mutagenesis, using the Q5 High-Fidelity 2X Master Mix protocol as well.

*Lf*ArsA plasmid was transformed into BL21 Gold (DE3) competent cells (Invitrogen). Starter overnight cultures (5-mL) were prepared by inoculated single colony in Luria broth media in the presence of 100 μg/mL ampicillin and grown for ∼18 h. Sterile 2xYT media (1 L) supplemented with 100 μg/mL ampicillin was inoculated with the overnight culture and grown at 37°C until OD reached ∼0.6, when the cells were induced with 0.4 mM isopropyl β-D-1-thiogalactopyranoside (IPTG), and grown at 18 °C for ∼16-18 h. Cells were harvested, flash frozen in liquid nitrogen and stored at −80°C. For purification, thawed cells were resuspended into ‘buffer A’ (50 mM HEPES pH 7.5, 300 mM NaCl, 10% glycerol, 20 mM imidazole, 5 mM β-mercaptoethanol (β-ME) and 0.2 mM phenylmethylsulfonyl fluoride (PMSF)), and lysed using an M-110L microfluidizer (Microfluidics). The debris was spun down at 24,000 xg for 30 min and the supernatant was incubated with Ni-NTA affinity resin (Qiagen) at 4°C for 1 h. Unbound material was removed by batch method using a bench-top centrifuge at 700 xg for 5 min, followed by 2x washes with 50 column-volumes of buffer A. At this point, the resin was transferred to an Econo-Pac chromatography column (Bio-Rad), and four fractions (1 column-volume each) were eluted with ‘buffer B’ (buffer A + 200 mM imidazole). After SDS-PAGE analysis, fractions containing ArsA were pooled, concentrated to 5 mL in an Amicon 30k concentration filter, and loaded onto a 120-mL HiLoad Superdex-200 column pre-equilibrated with 50 mM HEPES pH 7.5, 100 mM NaCl, 10% glycerol and 5 mM dithiothreitol (DTT). Alternatively, 5 mM tris(2-carboxyethyl)phosphine (TCEP) was used as reducing agent for storage buffer. After SDS-PAGE analysis, pure fractions were stored at −80°C. Protein concentration was determined by measuring the absorbance at 280 nm with a NanoDrop 2000 spectrophotometer (Thermo Fisher), using molar extinction coefficient of 25,400 M^−1^cm^−1^ and molecular weight of 63.8 kDa.

### X-ray absorption spectroscopy of ArsA

‘As^III^-ArsA’ sample was prepared by incubating 1.95 mM of *Lf*ArsA D46N/D364N (non-hydrolyzing variant) with 15 mM MgCl_2_, 15 mM ATP (Sigma), and 1.5 mM of sodium arsenite (Sigma) and then buffer exchanged into 50 mM HEPES pH 7.5, 100 mM NaCl, 30% glycerol and 5 mM TCEP using Micro Bio-Spin P-6 desalting column (Bio-Rad) to remove excess unbound arsenite. ‘As^III^-ArsA C173A’ sample was prepared similarly with 2.5 mM of *Lf*ArsA (non-hydrolyzing variant) C173A incubated with 20 mM MgCl_2_, 20 mM ATP, and 20 mM sodium arsenite. The ‘As^III^-cysteine’ sample (control) was prepared by adding 1.2 mM sodium arsenite to 6 mM L-cysteine in 50 mM HEPES pH 7.5, 100 mM NaCl, 30% glycerol, and 5 mM TCEP. A replicate was prepared and analyzed for each sample (Fig. S13 & Table S3). As^III^-ArsA replicate was supplemented with 15 mM sodium arsenite instead. Samples (∼150 µL each) were injected into leucite cells using a Hamilton syringe, avoiding air bubbles, and immediately frozen in liquid nitrogen.

XAS data were collected at the Stanford Synchrotron Radiation Laboratory (SSRL) on beamline 7-3, equipped with Si[220] double-crystal monochromator with a harmonic rejection mirror. Samples were maintained at 10 K using an Oxford Instruments continuous-flow liquid helium cryostat. Protein fluorescence excitation spectra were recorded using a 30-element Ge solid-state array detector. A germanium filter (0.6 mm in width) and solar slits were placed between the cryostat and detector to filter scattering fluorescence not associated with protein-bound arsenic signals. XAS spectra were recorded in 5 eV steps in the pre-edge region (11625– 11825 eV), 0.25 eV steps in the edge region (11850–11900 eV), and 0.05 Å^−1^ increments in the extended X-ray absorption fine structure (EXAFS) region out to a *k* range of 14 Å^−1^. The data were integrated from 2 to 25 s in a *k*-weighted manner in the EXAFS region for a total scan length of 45 min. X-ray energies were calibrated using an arsenic foil absorption spectrum collected simultaneously with the protein data. The first inflection point for the arsenic foil edge was assigned to 11867 eV. Each fluorescence channel of each scan was examined for spectral anomalies prior to averaging. The data represent an average of five to six scans for each sample. Data were processed using the Macintosh OS X version of the EXAFSPAK software suite integrated with Feff version 7 for theoretical model generation. Data, collected out to *k* = 14.0 Å^− 1^, corresponds to a spectral resolution of 0.121 Å^−1^ for all metal–ligand interactions; therefore, only independent scattering environments at distances >0.121 Å were considered resolvable in the EXAFS fitting analysis. The final EXAFS fitting analysis was performed on raw/unfiltered data. Protein EXAFS data were fit using single-scattering F_eff_ theoretical models calculated for carbon, oxygen, and sulfur coordination to simulate arsenic-ligand environments, with values for the scale factors (0.98) and E_0_ (−10) following a previously published fitting protocol. All spectra were fit using identical protocols, first by distinguishing the best single-shell fit to the data and then by progressively adding extra scattering environments to the fit. Best fit selection criteria were identified by having the lowest mean square deviation between experimental data and the theoretical fit (*F^’^* value), along with an acceptable absorber-scatterer bond disorder value (Debye-Waller factor) of < 6.0 × 10^3^ Å^2^.

### Cryo-EM sample preparation

For the *ADP-open* structure, the *Lf*ArsA sample was buffer-exchanged into 50 mM HEPES pH 7.5, 100 mM NaCl, and 4 mM TCEP using Micro Bio-Spin P-6 desalting columns (Bio-Rad). The resulting sample was diluted to 10 mg/mL and incubated with 2 mM each of MgCl_2_, ADP (Sigma), and 5 mM sodium arsenite for ∼3 hours. Samples for As^III^-free structures were prepared similarly. For the *ATP-closed* structure, *Lf*ArsA D46N/D364N was buffer-exchanged into 50 mM HEPES pH 7.5, 100 mM NaCl, and 5 mM DTT. The resulting sample was diluted to 10 mg/mL and incubated with 2 mM each of MgCl_2_, ATP, and sodium arsenite for ∼30 min. The sample for *wt*ArsA *ATP-closed* structure was prepared similarly but incubated with ligands for 1.5 min prior to grid preparation. For cryo-EM grid preparation, 3 μL of the sample supplemented with 0.05% CHAPSO was applied to glow-discharged Quantifoil holey carbon R1.2/1.3 300 Mesh, Copper (Quantifoil, Micro Tools GmbH) grids using a Vitrobot (FEI Vitrobot Mark v4 x2, Mark v3). Grids were blotted at 100% humidity and 4°C using a blot time of 4-5 seconds and blot force of 7, and immediately followed by plunge-freezing into liquid ethane.

### Cryo-EM data acquisition and processing

The grids were screened for ice thickness and sample quality using a 200 kV Talos Arctica TEM equipped with a Gatan K3 detector. Data collection was performed using a 300 kV Titan Krios TEM equipped with a Gatan K3 direct electron detector and Gatan Energy Filter (slit width 20eV) in super-resolution mode using SerialEM^43^. Each dataset was acquired at a nominal magnification of 130,000x with a raw pixel size of 0.325 Å/pixel, electron exposure of 70 e^−^/Å ^2^ over 40 frames (exposure rate of 1.75 e^−^/Å^2^/frame), and a defocus range of −0.5 to −2.5 μm. Correlated double sampling (CDS) mode was enabled to improve the signal-to-noise ratio of the images^44^.

All datasets were processed in cryoSPARC v4.4.1-4.5.1 using the same overall processing workflow^45^. Movie frames were motion corrected using ‘patch motion correction’ with 0.5 F-cropping, resulting in a pixel size of 0.65 Å/pixel. The contrast transfer function (CTF) for the motion-corrected micrographs was estimated using ‘patch CTF estimation’. Micrographs were manually curated, and particles were picked using ‘blob-picking’, extracted with a 2x bin (1.3 Å/pixel), and subjected to multiple rounds of 2D classification to retain good-quality particle picks. Ab-initio reconstruction was performed on this particle set to result in four 3D volumes, followed by ‘heterogenous refinement’ that yielded one good volume revealing secondary structural features of ArsA. Particles were re-extracted from the micrographs with no binning (0.65 Å/pixel) and subjected to multiple rounds of ab-initio reconstruction and heterogeneous refinement followed by ‘reference-based motion correction’, ‘global CTF refinement’, and ‘non-uniform refinement’^46^, to obtain a final set of particles resulting in a good-quality map. The overall resolution was estimated from the gold-standard Fourier shell correlation (FSC) curve at a cut-off of 0.143 in cryoSPARC. B-factor sharpening values were determined using ‘sharpening tools’ in cryoSPARC. Sharpened maps were exported from cryoSPARC for model building and refinement. The local resolution of each map was calculated using ‘local resolution estimation’ job in cryoSPARC. For the ArsA•MgATP map, iterative rounds of *ab-initio* reconstruction were performed instead to obtain the final set of particles. The image processing pipeline for each cryo-EM map reported in this work is presented in the SI Appendix, Fig. S3 (ArsA•MgADP in the presence of As^III^), Fig. S6 (ArsA•MgADP in the absence of As^III^), Fig. S7 (ArsA•MgATP), Fig. S8 (ArsA•MgATP•As^III^), and Fig. S11 (*wt*ArsA•MgADP•As^III^).

### Model building and refinement

All models were built and refined similarly unless stated otherwise. Initial models were obtained by docking the N (residues 1-294) and C (residues 316-587) domains of *Lf*ArsA AlphaFold2^35^ model prediction determined using ColabFold v1.5.2, as distinct model entries into the sharpened EM maps using ‘Dock in map’ in Phenix v1.21.2^47^. The model was adjusted by manual model building into good-quality regions of the map in Coot v0.9.8.7^48^. Ligands (Mg^2+^, nucleotides and/or As^III^) and any ordered water molecules in the nucleotide-binding sites were docked in Coot. Models were refined using ‘Real-space refinement’ in Phenix and ‘Real-space refine zone’ in Coot. Custom geometry restraints for As-S bond length were obtained from the XAS data and applied to the As^III^ binding site during refinement cycles in Phenix for the ArsA•MgATP•As^III^ structure. Each As-S bond length was restrained to 2.27 Å with a sigma-value of 0.06 Å derived from the Debye-Waller factor *σ*^2^ of 3.49 × 10^3^ Å^2^ corresponding to As-S bond distance of ‘As^III^-ArsA’ sample (Table S2). Average and per-residue Q-scores for each refined model were calculated using the Qscore plugin in ChimeraX v1.7.1.

### ATPase assays

Steady-state ATPase activity of ArsA was measured spectrophotometrically using an NADH-linked coupled assay with an ATP regeneration system that couples ATP hydrolysis to oxidation of NADH^49^. The reaction mixture (100 µL) consisted of 50 mM HEPES pH 7.5, containing 5 mM MgCl_2_, 2 mM phosphoenol pyruvate (Roche), 20 U/mL pyruvate kinase from rabbit muscle (MP Biomedicals), 20 U/mL L-lactate dehydrogenase from rabbit muscle (Sigma), 0.2 mM NADH (Roche), 2-5 µM ArsA or its variant and varying concentrations of sodium arsenite (0, 0.05, 0.1, 0.2, 0.5 and 0.8 mM). The mixture was incubated for 5-10 min at 37 °C. The assay was conducted in a 96-well plate setup using SpectraMax M3 plate reader (Molecular Devices). The reaction was initiated by adding 5 mM ATP into the wells with gentle mixing. This was immediately followed by the measurement of a steady-state decrease in the NADH absorbance at 340 nm for 20 min on the plate reader at 37°C. Slopes were calculated for the steady region of the progress curves in the units of A_340_ min^−1^. Rates were calculated as nanomoles of ATP hydrolyzed per min per mg of ArsA (nmol min^−1^ mg^−1^) using the extinction coefficient of NADH at 340 nm (6,220 M^−1^ cm^−1^). ATPase activity was fitted to the Michaelis-Menten equation as a function of arsenite concentration and plotted using Python 3 in Jupyter notebook with assistance from ChatGPT, OpenAI in writing the code.

### Inductively coupled plasma mass spectroscopy (ICP-MS)

ArsA-bound arsenic was quantified for ArsA•MgATP, ArsA C173A•MgATP, ArsA•MgADP and ArsA C114A/C173A/C422A•MgADP samples. The non-hydrolyzing variant was utilized when trapping complexes with MgATP. Each sample was prepared similar to the cryo-EM samples by incubating 10 mg/mL of *Lf*ArsA or its variant, with 2 mM MgCl_2,_ 2 mM ATP/ADP, and 5 mM sodium arsenite at 4°C for 3 hours. Free arsenite was removed from the samples by exchanging the buffer with 50 mM HEPES pH 7.5 using Micro Bio-Spin P-6 desalting column. Subsequently, arsenic was extracted from the protein by incubating 30 µL of the sample with 1.1 mL of 70% (v/v) HNO_3_ (ACS grade) in borosilicate glass tubes at ∼50°C for 30 min. Each sample was then diluted to 15 mL with double distilled water in a 50-mL flat bottom tube, resulting in maximum arsenic concentration equivalent to 23.5 ppb or 0.313 µM of ArsA in 5% (v/v) HNO_3_.

Arsenic concentrations were determined by ICP-MS using an Agilent 8800. The sample introduction system consisted of a Miramist nebulizer, Scott-type spray chamber, and 2.0 mm fixed injector quartz torch. A guard electrode was used, and the plasma was operated at 1500 W. Arsenic analysis was performed in He mode using MS/MS scan mode. Arsenic standards were prepared in 5% (v/v) HNO_3_ from a 10 µg/mL arsenic standard solution (Inorganic Ventures) in the range of 0 to 40 ppb of As. The probe was subjected to 4 rinses (1 flowing, 3 static) between samples to minimize any possibility of cross contamination. Sample to sample carryover has been found to be less than 1% between the highest standard (40 ppb) and the blank for similar analyses. Results were analyzed using ICP Masshunter 4.5 (Agilent Technologies).

## Data Availability

Cryo-EM maps and the corresponding atomic models generated in this study have been deposited into the Protein Data Bank (PDB) and the Electron Microscopy Data Bank (EMDB) for release upon publication. Scripts for the ATPase assay and ICP-MS plots can be accessed at https://github.com/smahajan82/ArsAplots.git.

## Acknowledgements

This work was supported by funding from Howard Hughes Medical Institute (D.C.R.), Chan Zuckerberg Initiative (W.M.C.), and the Center for Environmental-Microbial Interactions at Caltech (S.M.). Cryo-EM data were collected at the Caltech Cryo-EM Resource Center supported by the Beckman Institute, and we are grateful to Songye Chen, Tyler J. Brittain, and Victor Garcia for their assistance with data collection and processing. XAS data was collected on beamline 7-3 at the Stanford Synchrotron Radiation Laboratory (SSRL), SLAC National Accelerator Laboratory which is supported by the U.S. Department of Energy and by the National Institutes of Health, National Institute of General Medical Science (P30GM133894). We thank the SSRL staff as well as Jens T. Kaiser for his assistance with preparing and shipping XAS samples. ICP-MS data was collected at the Caltech Water Exploration Laboratory (WEL) supported by the Resnick Sustainability Institute, and we thank Nathan F. Dalleska for assistance with running ICP-MS and analyzing data. We are grateful for access to the SpectraMax M3 plate reader for ATPase assays in the Dianne K. Newman lab at Caltech. Finally, we are grateful to Xiang Feng, Ailiena O. Maggiolo, Rebeccah A. Warmack, Jacob M. Kirsh, and Juliet A. Lee for discussions and critical reading of the manuscript.

## Author contributions

S.M., D.C.R. and W.M.C. designed research; S.M., Y.E.L., A.E.P. and T.L.S. performed research; S.M., T.L.S., D.C.R. and W.M.C. analyzed data; S.M., D.C.R. and W.M.C. wrote the manuscript.

