## supplementary_information for "Nucleotide and metalloid-driven conformational changes in the arsenite efflux ATPase ArsA"

[1]Present address for Timothy L. Stemmler: Medical University of South Carolina, Charleston, SC 29425.

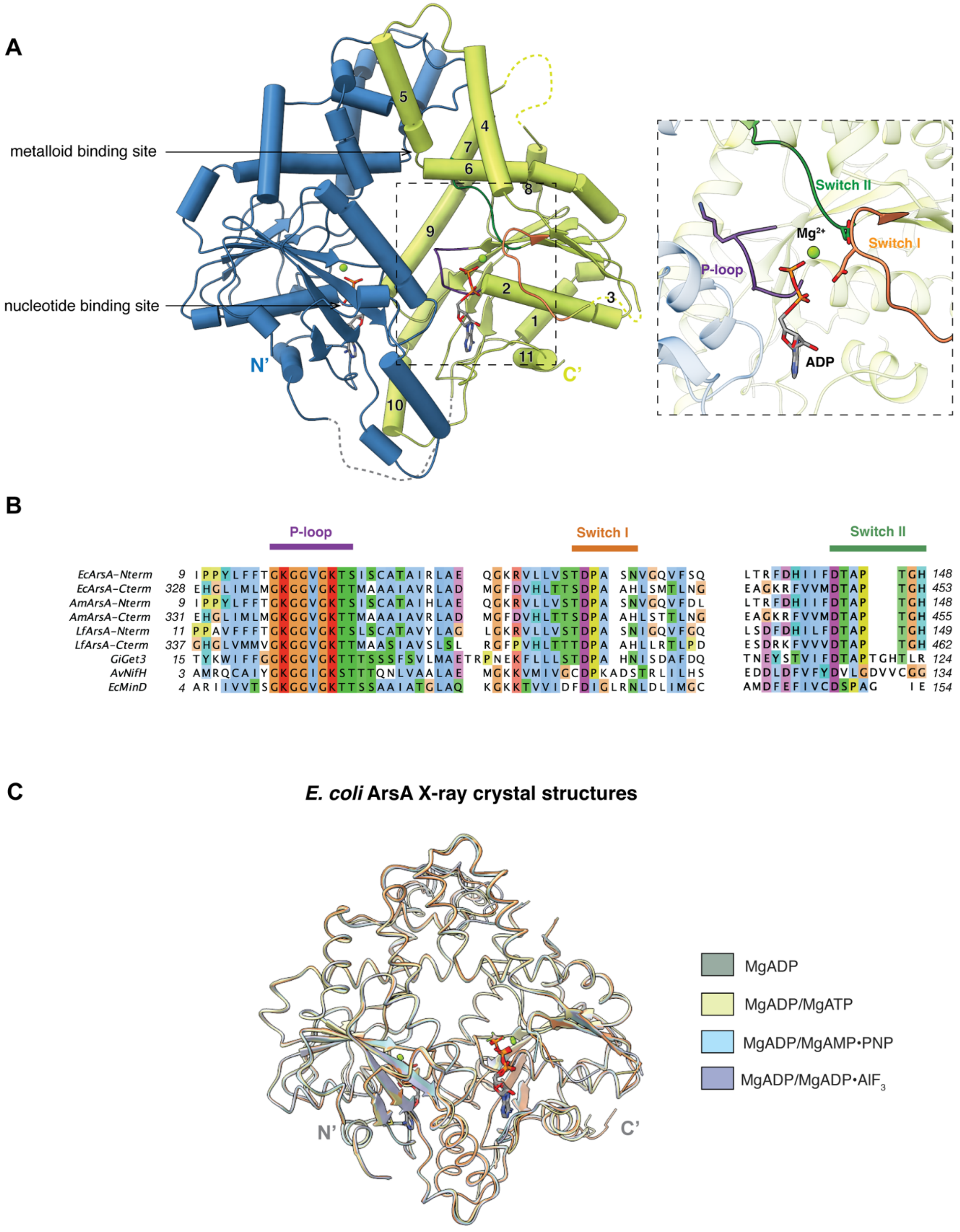


**Fig. S1. Arsenite efflux ATPase ArsA from *E. coli*. (A)** *Ec*ArsA structure bound to MgADP (PDB: 1F48) highlighting the N- (*blue*) and C- (*green*) domains, the nucleotide-binding sites and the metalloid-binding site. The inset highlights the P-loop (violet), Switch I (orange) and Switch II (green) motifs of the C-domain. Helices are numbered 1 through 11 on the C-domain, and have homologous helices on the N-domain. Helix 3 is not modeled in the C-domain of this structure, but the N-domain helix 3 is shown. **(B)** Multiple sequence alignment of N- and C-domains of ArsA homologues from *E. coli* (*Ec*ArsA), *Acidiphilium multivorum* (*Am*ArsA), and *Leptospirillum ferriphilum* (*Lf*ArsA) with other IWA ATPases – Get3 from *Giardia intestinalis* (*Gi*Get3), Nitrogenase Fe protein from *Azotobacter vinelandii* (*Av*NifH), and bacterial cell division regulatory ATPase from *E. coli* (*Ec*MinD) highlighting the conserved IWA motifs as shown in *A*. Alignment was constructed using Promals3D[1] and residues were colored using the ClustalX scheme[2]. **(C)** Global alignment of all X-ray crystal structures of *Ec*ArsA bound to various nucleotide – MgADP (PDB: 1F48), MgADP/MgATP (PDB: 1II0), MgADP/MgAMP•PNP (PDB: 1II9), and MgADP/MgADP•$AlF_3$ (PDB: 1IHU). MgADP is bound at the N-domain and a different nucleotide is bound at the C-domain in each structure. $Sb^{III}$/$Cd^{2+}$ and $Cl^-$ ions are modeled at the metalloid-binding sites in each of these structures but were omitted from this representation.

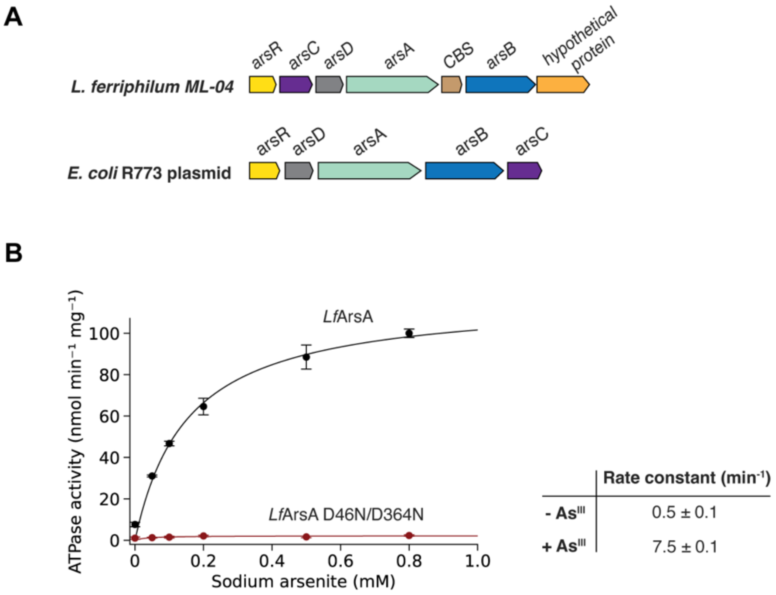


**Fig. S2. Arsenite efflux pump ATPase ArsA from *Leptospirillum ferriphilum* strain ML-04. (A)** Architecture of the *ars* operon in *L. ferriphilum* ML-04 in comparison with that of the *E. coli* plasmid R773. Both operons constitute genes for an $As^{III}$-responsive repressor, arsR; $As^{III}$-metallochaperone, arsD; $As^{III}$-translocating ATPase, arsA; the membrane transporter, arsB; and arsenate ($As^{V}$) reductase, arsC. The *L. ferriphilum ars* operon consists of two additional genes – a cystathione-β-synthase (CBS) domain regulatory protein, and a hypothetical protein annotated as a putative periplasmic component of an ABC-type phosphate transporter[3]. **(B)** Steady-state ATPase activity of *Lf*ArsA (black plot) and the Switch I aspartate double mutant, D46N/D364N (red plot) as a function of $As^{III}$ concentration with 5 mM each of $MgCl_2$ and ATP at 37ºC. While the mutant is non-hydrolyzing, wild-type ArsA shows $As^{III}$-dependent activation of ATPase activity. In the absence of $As^{III}$ (0 mM data point), the calculated pseudo first-order rate constant is 0.5 ± 0.1 $min^{-1}$ (- $As^{III}$) and saturating $As^{III}$ concentrations enhance the rate constant to 7.5 ± 0.2 $min^{-1}$ (+ $As^{III}$). Data points represent mean of n = 3 and error bars represent standard deviations. The data was fit to the Michaelis-Menten equation. Rate constants were calculated using ArsA molecular weight of 63.8 kDa.

### Table S1. Cryo-EM data collection, refinement and validation statistics.

| | *Lf*ArsA•MgADP (+ $As^{III}$) (EMD-492321) (PDB: 9NBL) | *Lf*ArsA•MgADP (- $As^{III}$) (EMD-49232) (PDB: 9NBM) | *Lf*ArsA•MgATP (EMD-49233) | *Lf*ArsA•MgATP•$As^{III}$ (D46N/D364N) (EMDB-49234) (PDB: 9NBO) | *Lf*ArsA•MgATP•$As^{III}$ (wild-type) (EMDB-49237) (PDB: 9NBW) |
|---|---|---|---|---|---|
| **Data collection and processing** | | | | | |
| Magnification | 130,000 | 130,000 | 130,000 | 130,000 | 130,000 |
| Voltage (kV) | 300 | 300 | 300 | 300 | 300 |
| Electron exposure ($e^{-}/Å^2$) | 70 | 70 | 70 | 70 | 70 |
| Defocus range (μm) | -0.5 to -2.5 | -0.5 to -2.5 | -0.5 to -2.5 | -0.5 to -2.5 | -0.5 to -2.5 |
| Pixel size (Å) (Super-resolution mode) | 0.325 | 0.325 | 0.325 | 0.325 | 0.325 |
| Movies | 4,732 | 2,122 | 3,994 | 4,031 | 4,509 |
| Total extracted particles | 2,643,084 | 1,824,091 | 1,673,570 | 4,265,167 | 1,449,927 |
| Final particles | 103,258 | 34,966 | 48,382 | 91,645 | 124,599 |
| Symmetry imposed | C1 | C1 | C1 | C1 | C1 |
| Map resolution (Å) (FSC 0.143 cut-off) | 3.4 | 3.8 | 6.6 | 3.0 | 3.0 |
| **Refinement** | | | | | |
| Initial model used | *Lf*ArsA AlphaFold model | *Lf*ArsA AlphaFold model | | *Lf*ArsA AlphaFold model | *Lf*ArsA AlphaFold model |
| Model composition: | | | | | |
| Protein residues | 558 | 552 | | 571 | 570 |
| Ligands | ADP: 2, MG: 2 | ADP: 2, MG: 2 | | ATP: 2, MG: 2, ARS: 1, HOH: 5 | ATP: 2, MG: 2, ARS: 1, HOH: 13 |
| Map sharpening *B* factor ($Å^2$) | -165.4 | -163.0 | | -124.1 | -133.0 |
| R.m.s. deviations: | | | | | |
| Bond lengths (Å) | 0.003 | 0.003 | | 0.004 | 0.002 |
| Bond angles (°) | 0.616 | 0.661 | | 1.024 | 0.561 |
| **Validation** | | | | | |
| MolProbity score | 1.63 | 1.41 | | 1.34 | 1.21 |
| Clashscore | 13.30 | 7.42 | | 6.25 | 4.32 |
| Poor rotamers (%) | 0 | 0 | | 0 | 0 |
| Ramachandran plot: | | | | | |
| Favored (%) | 98.18 | 98.16 | | 98.05 | 98.23 |
| Allowed (%) | 1.82 | 1.84 | | 1.95 | 1.77 |
| Outliers (%) | 0 | 0 | | 0 | 0 |

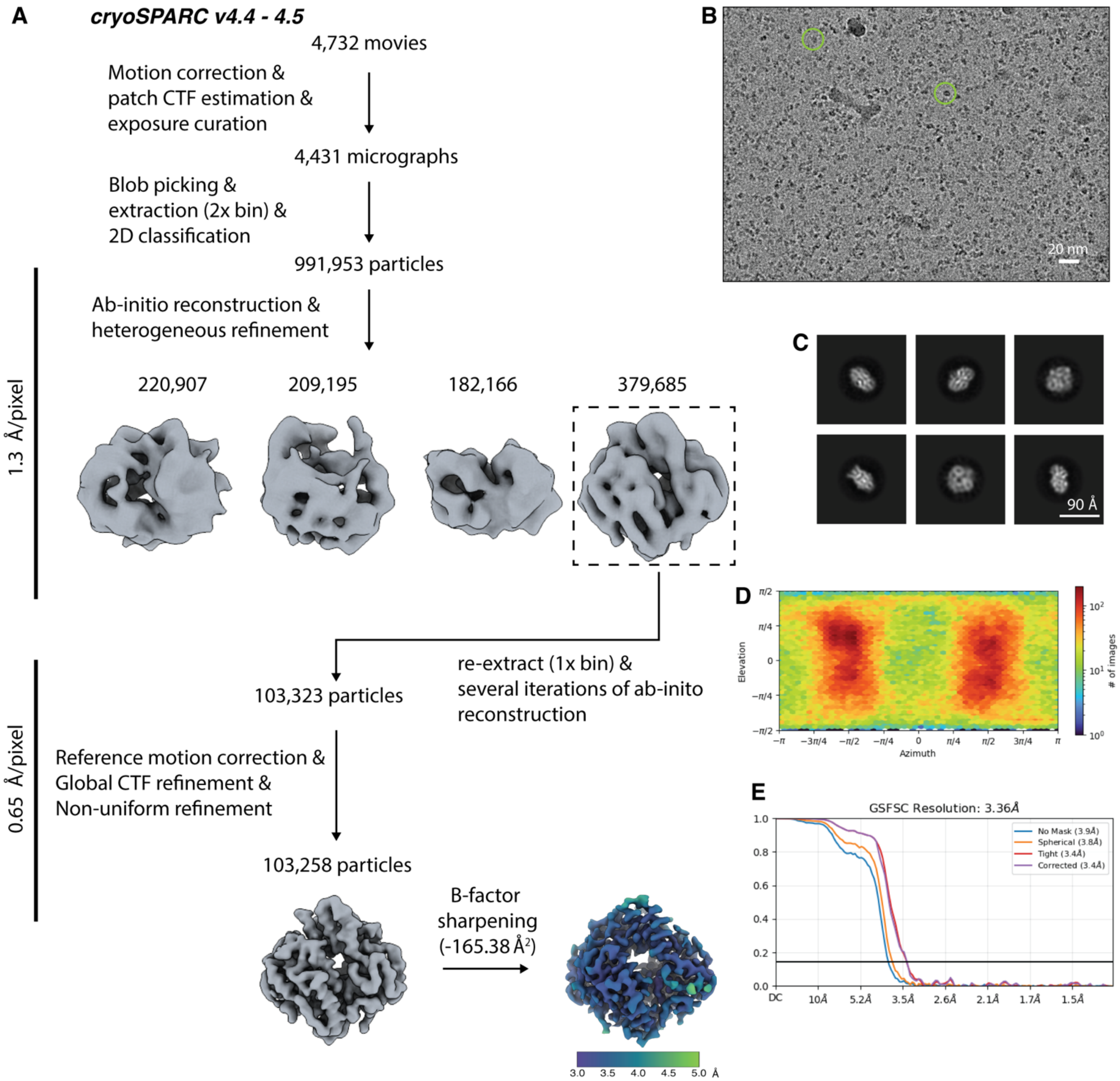


**Fig. S3. Cryo-EM processing and validation for *Lf*ArsA•MgADP structure determined in presence of As$^{III}$. (A)** Cryo-EM data processing workflow in cryoSPARC. **(B)** Representative micrograph. **(C)** 2D class averages showing different views of ArsA in the *open* state. **(D)** Angular distribution heatmap plot. **(E)** Gold-standard Fourier shell correlation (GSFSC) curve.

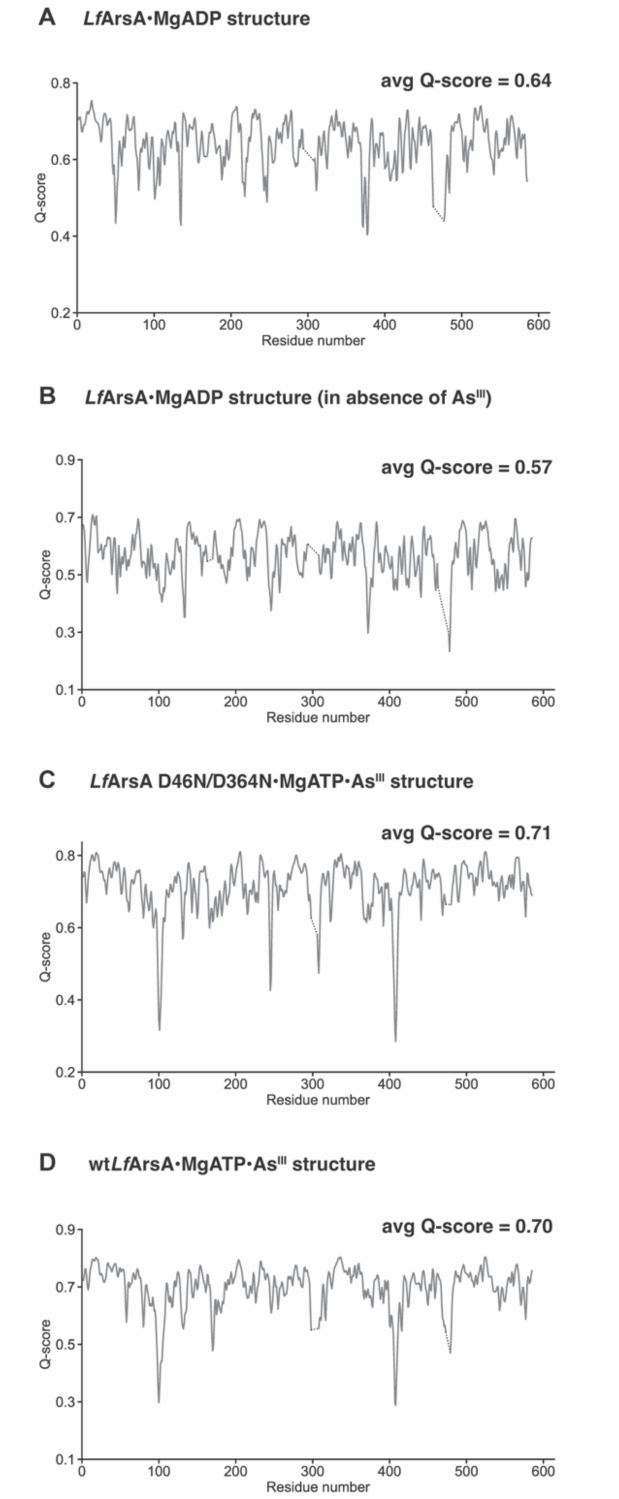


**Fig. S4. Per-residue average Q-scores for ArsA cryo-EM structures. (A)** *Lf*ArsA•MgADP (in presence of $As^{III}$), **(B)** *Lf*ArsA•MgADP (in absence of $As^{III}$), **(C)** *Lf*ArsA D46N/D364N•MgATP•$As^{III}$ and **(D)** wild-type *Lf*ArsA•MgATP•$As^{III}$. Average Q-scores are reported for residues 2-585 and were calculated using the Qscore plugin in ChimeraX v1.7.1. Unmodeled residues are shown as dotted lines.

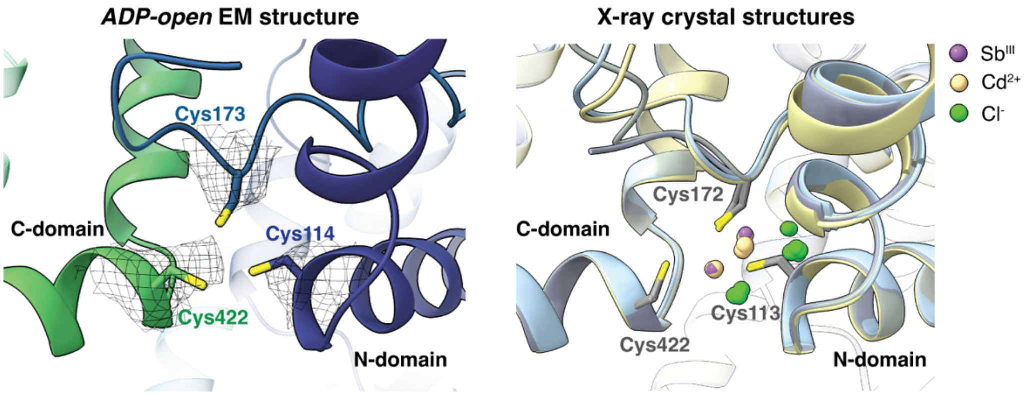


**Figure S5. Comparison of the metalloid-binding sites of *Lf*ArsA *ADP-open* cryo-EM structure (left) and *Ec*ArsA X-ray crystal structures (right).** While the latter includes $Sb^{III}$, $Cd^{2+}$ and $Cl^{-}$ ions modeled at the metalloid-binding site, the relative positions of the Cys residues are similar in both EM and X-ray structures. Likely, the ions present in the crystal structures were non-specifically acquired from the crystallization conditions, and therefore, do not represent a functionally relevant metalloid-bound state of ArsA. Crystal structures shown on the right include MgADP-bound (*gray*, PDB: 1F48), MgADP/MgATP-bound (*yellow*, PDB: 1II0), MgADP/MgAMP•PNP (*blue*, PDB: 1II9) and MgADP/MgADP•$AlF_3$ (*purple*, PDB: 1IHU) states.

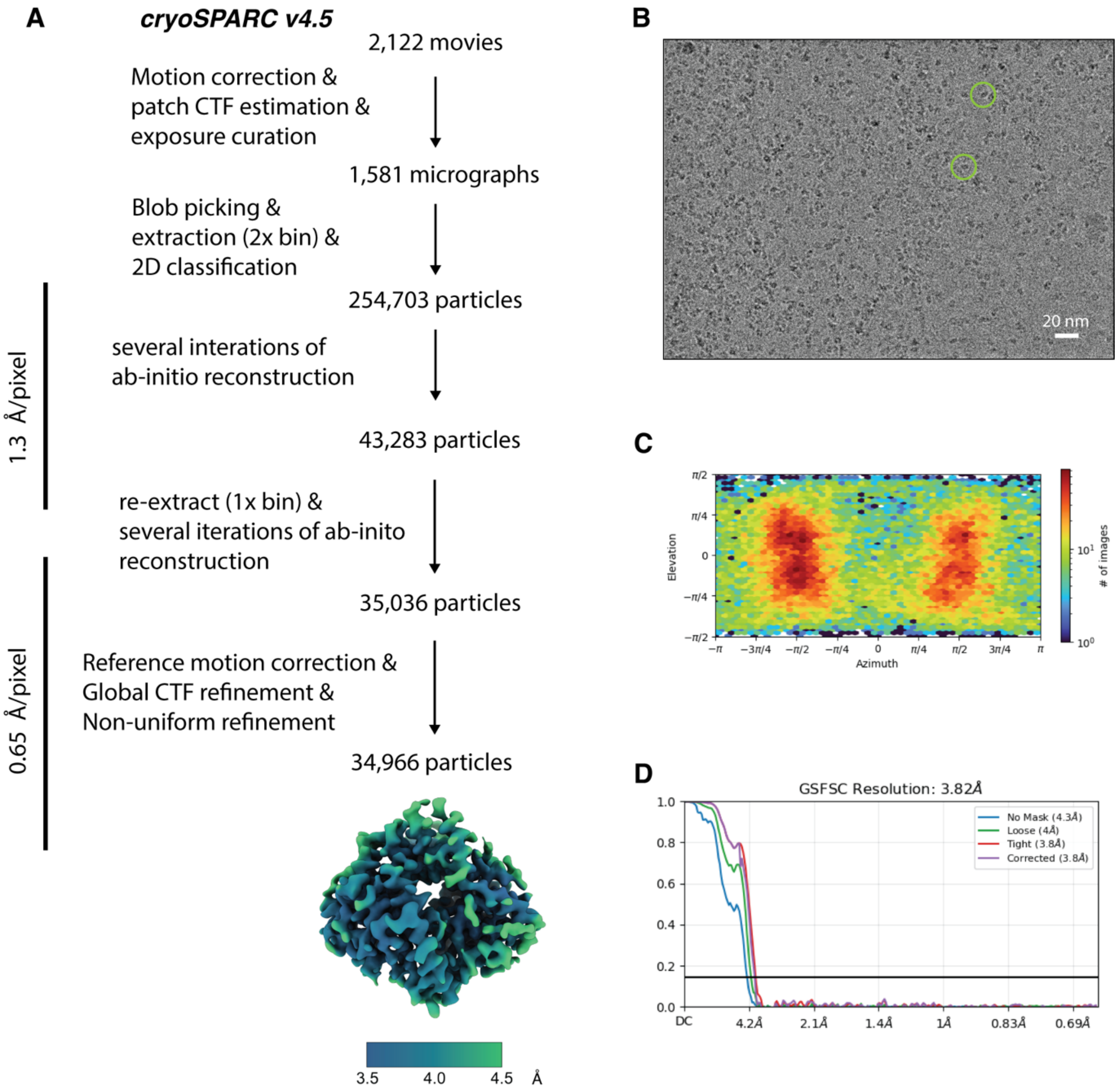


**Fig. S6. Cryo-EM processing and validation for *Lf*ArsA•MgADP structure determined in absence of As$^{III}$. (A)** Cryo-EM data processing workflow in cryoSPARC. **(B)** Representative micrograph. **(C)** Angular distribution heatmap plot. **(D)** Gold-standard Fourier shell correlation (GSFSC) curve.

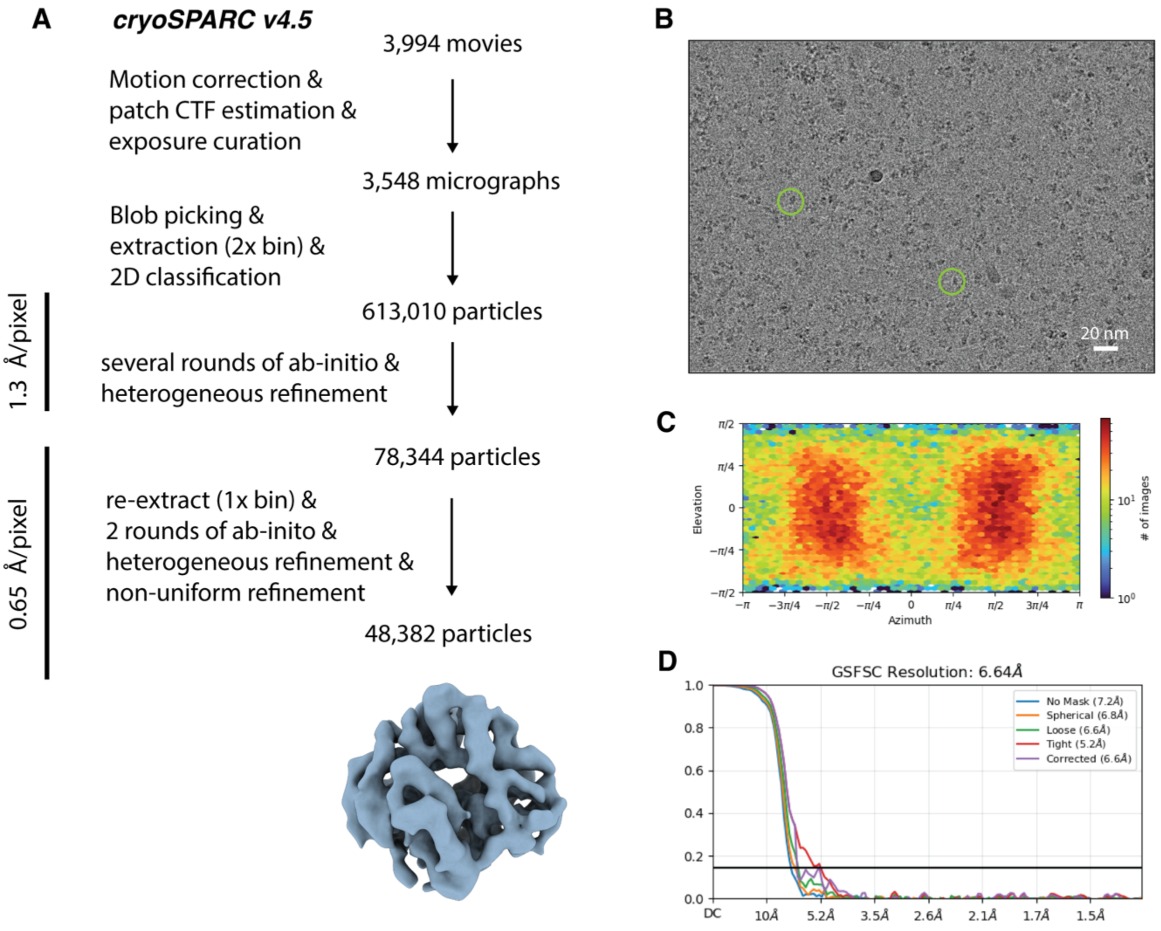


**Fig. S7. Cryo-EM processing and validation for *Lf*ArsA•MgATP structure. (A)** Cryo-EM data processing workflow in cryoSPARC. **(B)** Representative micrograph. **(C)** Angular distribution heatmap plot. **(D)** Gold-standard Fourier shell correlation (GSFSC) curve.

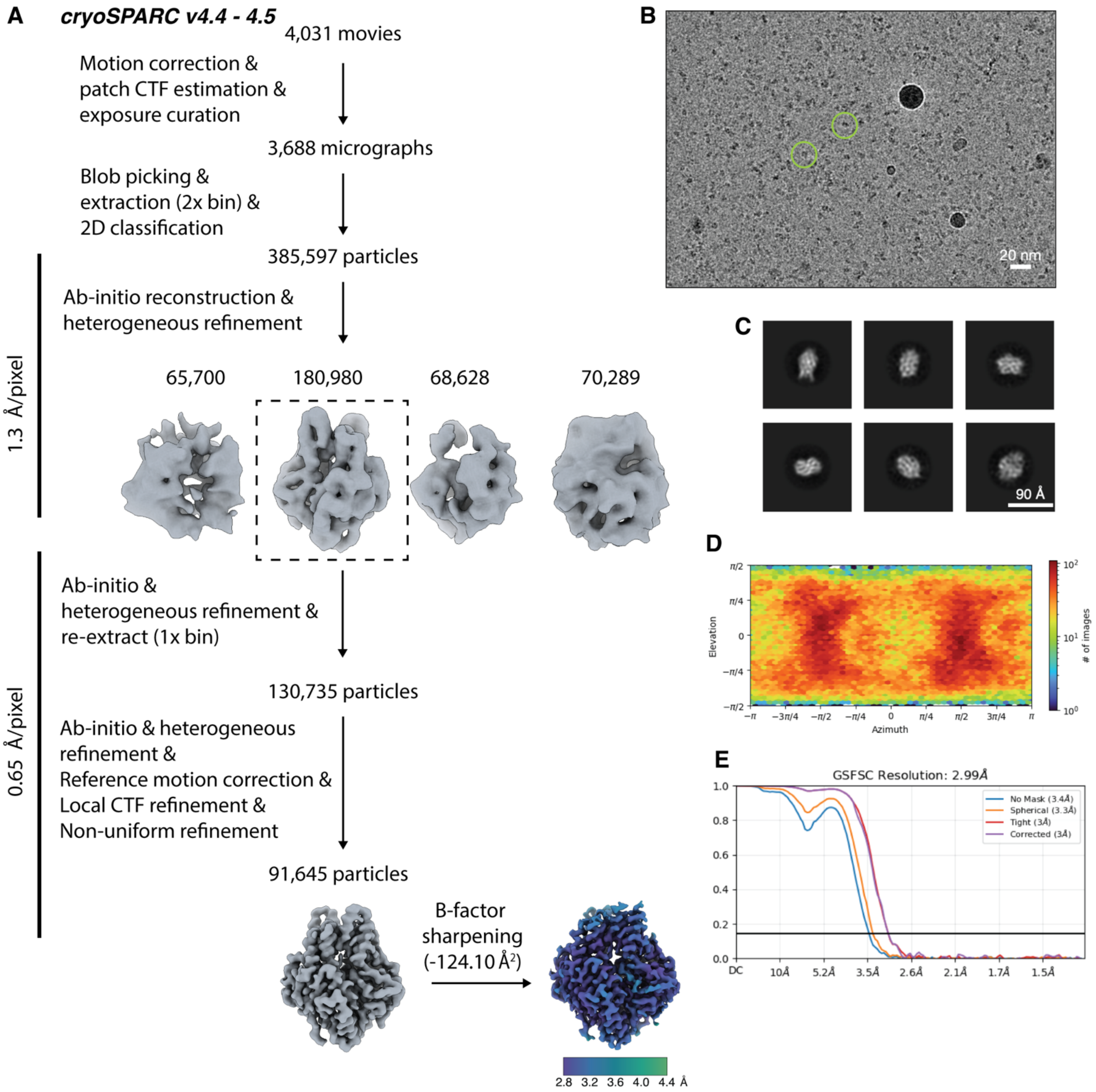


**Fig. S8. Cryo-EM processing and validation for *Lf*ArsA•MgATP•As$^{III}$ (non-hydrolyzing variant) structure. (A)** Cryo-EM data processing workflow in cryoSPARC. **(B)** Representative micrograph. **(C)** 2D class averages showing different views of ArsA in the *closed* state. **(D)** Angular distribution heatmap plot. **(E)** Gold-standard Fourier shell correlation (GSFSC) curve.

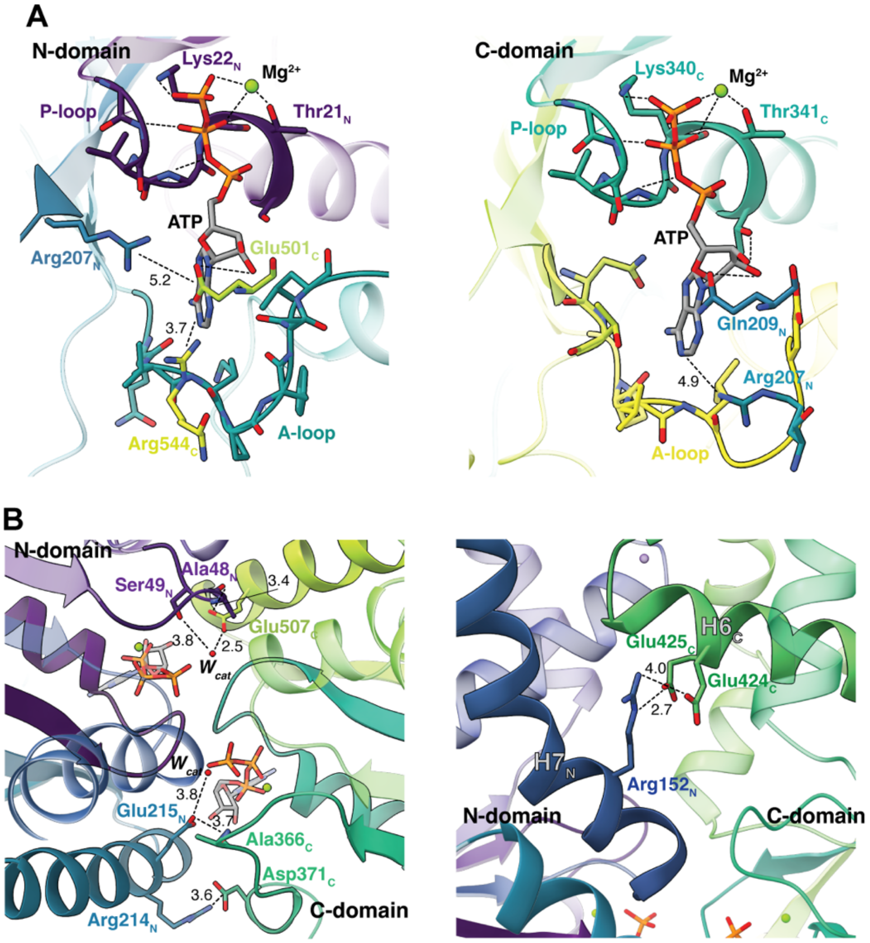


**Fig. S9. Residue interactions observed in the *ATP-closed* state of *Lf*ArsA. (A)** Residues from both domains interacting with ATP bound at N- (left) and C-sites (right). **(B)** Inter-domain contacts formed at the pseudodimer interface. Left, top-down view and right, front view of the pseudodimer showing interactions between N- and C-domains. All interacting residues are shown as sticks. The structure referred to here is from the non-hydrolyzing ArsA variant colored in viridis.

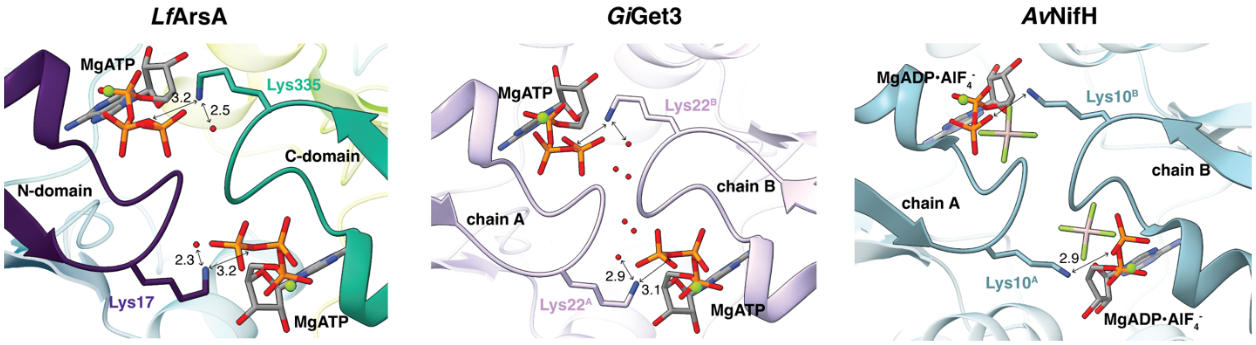


**Fig. S10. Comparison of P-loop interactions at the pseudodimer interface of the *ATP-closed* state of *Lf*ArsA (left) with that of *Gi*Get3 (center) and *Av*NifH (right) *closed* dimer state structures.** Waters at the interface are shown as *red* spheres (*Gi* – *Giardia Intestinalis* and *Av* – *Azotobacter vinelandii*).

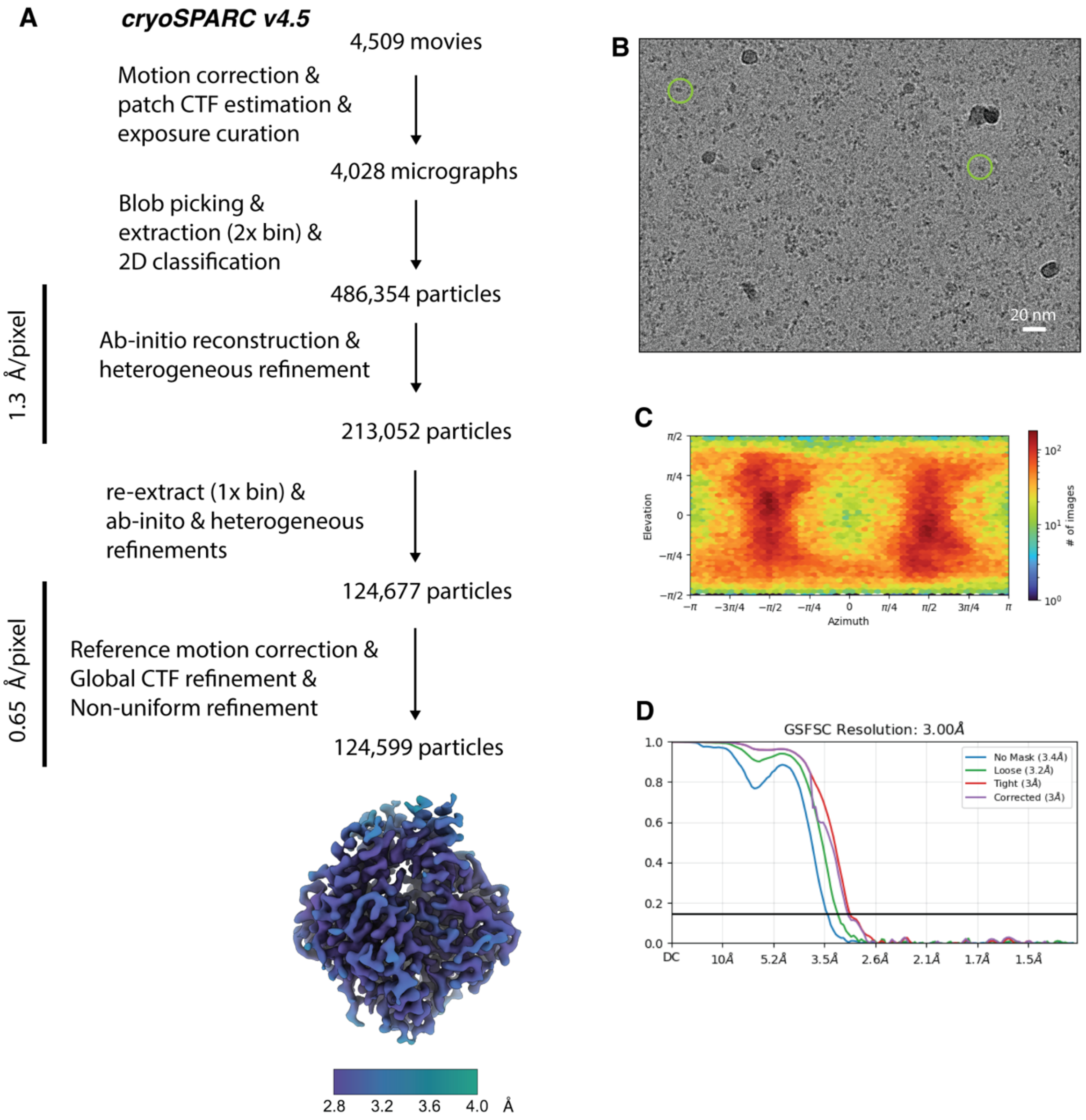


**Fig. S11. Cryo-EM processing and validation for *Lf*ArsA•MgATP•As$^{III}$ (wild-type) structure. (A)** Cryo-EM data processing workflow in cryoSPARC. **(B)** Representative micrograph. **(C)** Angular distribution heatmap plot. **(D)** Gold-standard Fourier shell correlation (GSFSC) curve.

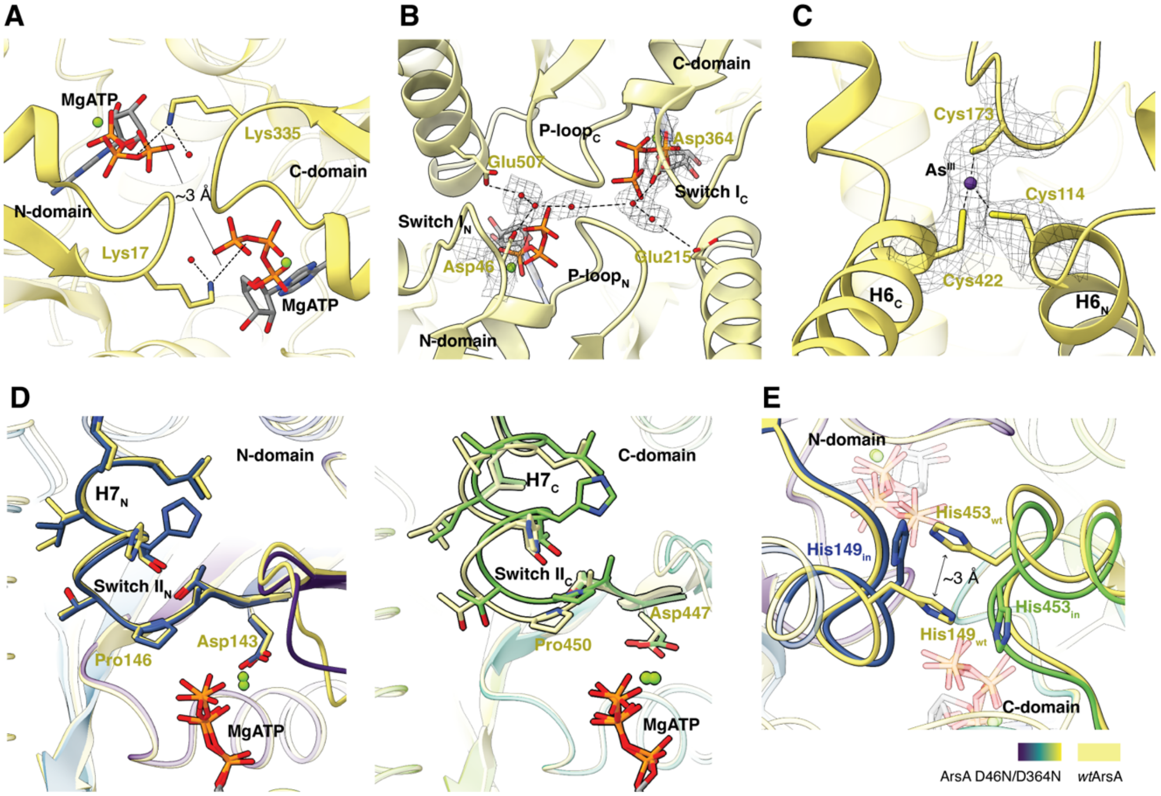


**Fig. S12. Structural features observed in the *ATP-closed* state of wild-type *Lf*ArsA. (A)** Pseudodimer interface showing that the P-loops of the two domains are in proximity and the IWA lysines (Lys17 and Lys335) are both oriented towards the ATP bound to the opposite domain. **(B)** Hydrogen-bonding network of water molecules (red) and Asp/Glu residues connecting the N- and C-sites. **(C)** Metalloid-binding site showing Coulomb potential map for As$^{III}$ (purple) enclosed between Cys114, Cys173 and Cys422. **(D)** Comparison of Switch II conformation between *wt*ArsA (*yellow*) and non-hydrolyzing variant (viridis) *closed* structures in the N- (left) and C- (right) domains. **(E)** His149 and His453 on helix 7 of N- and C-domains respectively, form a stacking interaction in the *wt*ArsA structure that is not observed in the non-hydrolyzing variant structure.

**Table S2. Arsenic EXAFS best fit simulation parameters.**

| Sample | Nearest Neighbor Ligand Environment[a] | | | | Long-Range Ligand Environment[b] | | | | |
|---|---|---|---|---|---|---|---|---|---|
| | Atom[c] | R(Å)[d] | C.N.[e] | $\sigma^2$ [f] | Atom[c] | R(Å)[d] | C.N.[e] | $\sigma^2$ [f] | *F'* [g] |
| $As^{III}$-ArsA | S | 2.27 | 3 | 3.49 | C | 2.74 | 1 | 3.39 | 0.66 |
| | | | | | C | 3.19 | 2 | 1.13 | |
| | | | | | C | 3.36 | 3 | 3.59 | |
| $As^{III}$-Cys | S | 2.27 | 2 | 3.06 | C | 3.21 | 1.5 | 4.84 | 0.30 |
| | | | | | C | 3.67 | 1 | 2.07 | |
| $As^{III}$-ArsA C173A | O/N | 1.78 | 2 | 3.80 | C | 2.72 | 2 | 3.90 | 0.34 |
| | S | 2.28 | 1 | 3.71 | C | 2.86 | 1.5 | 3.85 | |

[a]/[b]- Independent metal-ligand scattering environment.
[c] - Scattering atoms: N (nitrogen), O (oxygen), C (carbon), S (sulfur).
[d] - Average metal-ligand bond length.
[e] - Average metal-ligand coordination number.
[f] - Average Debye-Waller factor ($Å^2 \times 10^3$).
[g] - Number of degrees of freedom weighted mean square deviation between data and fit.

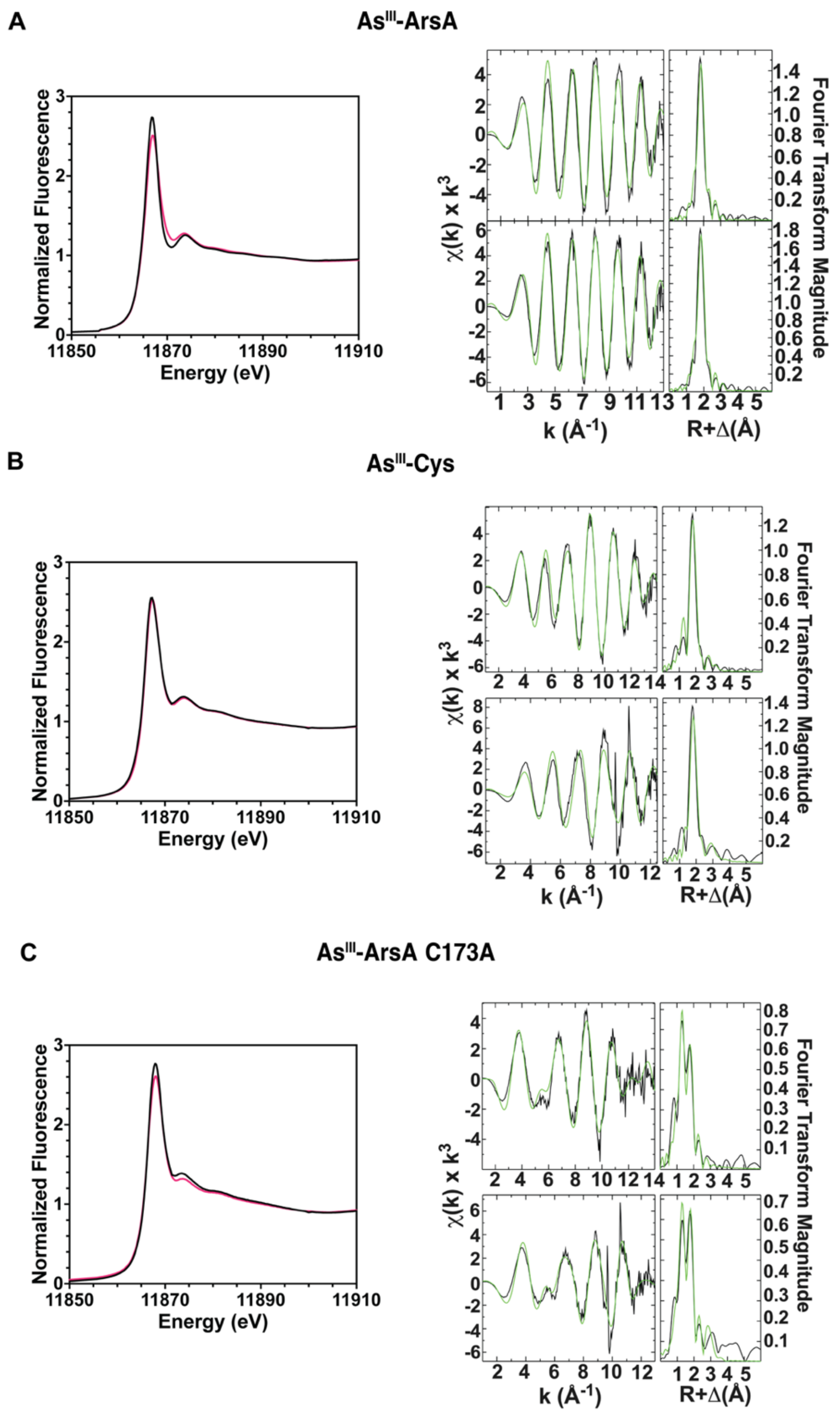


**Fig. S13. Duplicate XAS data for ArsA samples.** XANES (left) and EXAFS (right) spectra for **(A)** *Lf*ArsA•MgATP•As$^{III}$ (As$^{III}$-ArsA), **(B)** As$^{III}$-L-cysteine control (As$^{III}$-Cys) and **(C)** *Lf*ArsA(C173A)•MgATP•As$^{III}$ (As$^{III}$-ArsA C173A). Duplicate XANES spectra are shown in black and red.

**Table S3. Arsenic EXAFS best fit simulation parameters for a replicate of each sample.**

| Sample | Nearest Neighbor Ligand Environment[a] | | | | Long-Range Ligand Environment[b] | | | | |
|---|---|---|---|---|---|---|---|---|---|
| | Atom[c] | R(Å)[d] | C.N.[e] | $\sigma^2$ [f] | Atom[c] | R(Å)[d] | C.N.[e] | $\sigma^2$ [f] | *F'* [g] |
| $As^{III}$-ArsA | S | 2.27 | 2.5 | 3.46 | C | 2.72 | 1 | 3.64 | 0.60 |
| | | | | | C | 3.20 | 2 | 1.81 | |
| | | | | | C | 3.36 | 3 | 5.47 | |
| $As^{III}$-Cys | S | 2.27 | 2 | 3.12 | C | 3.31 | 2 | 3.08 | 1.28 |
| | | | | | C | 3.63 | 1.5 | 2.23 | |
| $As^{III}$-ArsA C173A | O/N | 1.78 | 2 | 4.06 | C | 2.71 | 2 | 3.11 | 0.61 |
| | S | 2.28 | 1 | 3.20 | C | 2.89 | 1 | 5.82 | |
| | | | | | C | 3.33 | 2 | 1.48 | |

[a]/[b]- Independent metal-ligand scattering environment.
[c] - Scattering atoms: N (nitrogen), O (oxygen), C (carbon), S (sulfur).
[d] - Average metal-ligand bond length.
[e] - Average metal-ligand coordination number.
[f] - Average Debye-Waller factor ($Å^2$ x $10^3$).
[g] - Number of degrees of freedom weighted mean square deviation between data and fit.

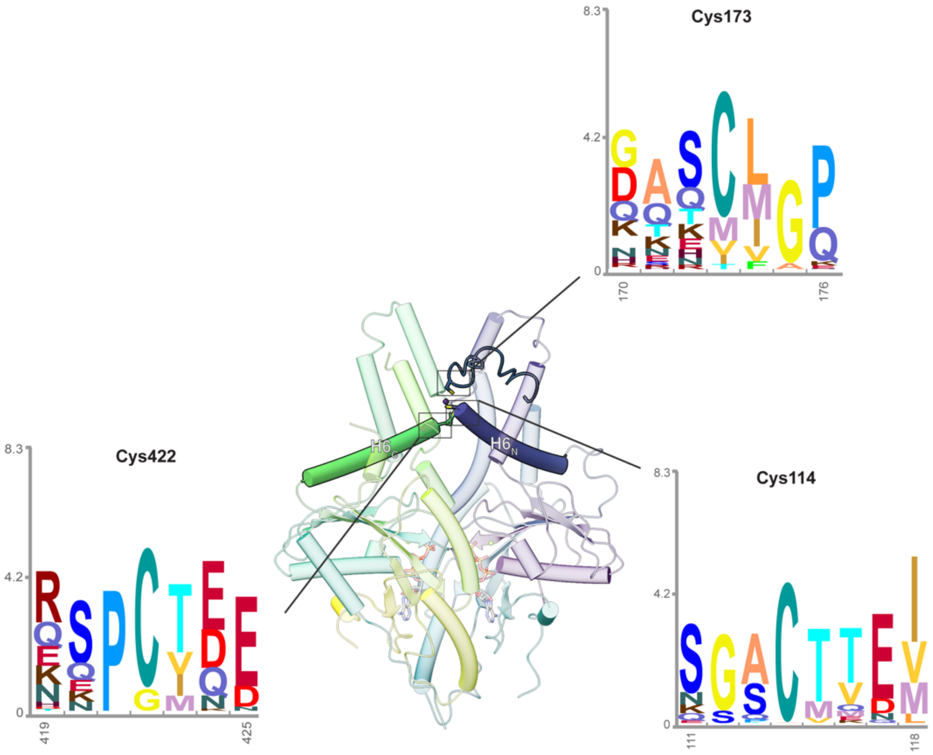


**Fig. S14. Conservation of Cys residues of the metalloid-binding site across ArsA homologs.** Hidden Markov Model (HMM) logos depict conservation of Cys114, Cys173 and Cys422 necessary for $As^{III}$ binding. Logos were constructed using 'Skylign' web tool[4] from a multiple sequence alignment of sequences assigned to the ArsA InterPro family (IPR027541).

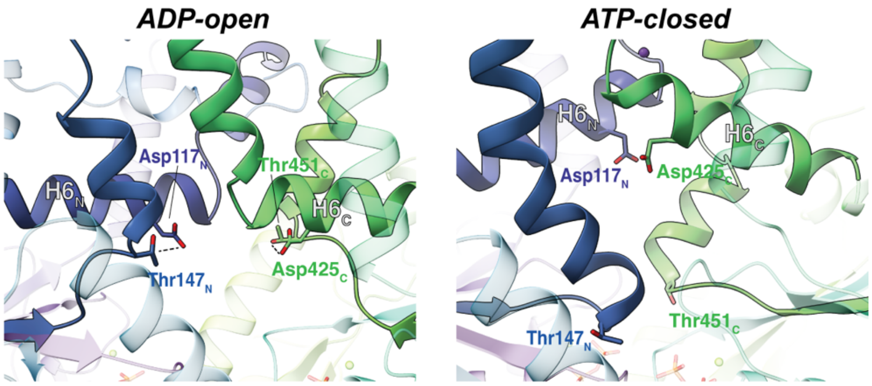


**Fig. S15. Interaction between Thr residue (Switch II) and Asp residue (helix 6) observed in the *ADP-open* (left) state is destabilized in the *ATP-closed* (right) state as Switch II and helix 6 undergo conformational changes in both N- and C-domains.**

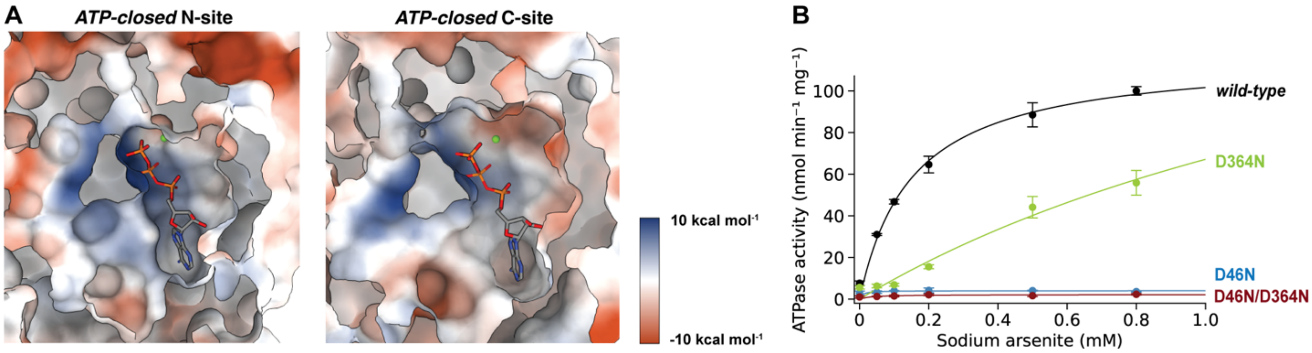


**Fig. S16. Non-equivalence of N- and C-domains of ArsA. (A)** Electrostatic potential surface representation of N- and C-sites of *Lf*ArsA *ATP-closed* state, showing that ATP binds in a tighter active site pocket in the N-site than in the C-site. Surfaces were generated in ChimeraX using the '*coulombic*' command. **(B)** Steady-state ATPase activity of ArsA (nmol $min^{-1}$ $mg^{-1}$) plotted against varying $As^{III}$ concentrations (mM), with 5 mM each of $MgCl_2$ and ATP at 37ºC (wild-type, black; D46N, blue; D364N, green and D46N/D364N, red). Data points represent mean of n = 3 and error bars represent standard deviations. The data was fit to the Michaelis-Menten equation.
